# SpikeShip: A method for fast, unsupervised discovery of high-dimensional neural spiking patterns

**DOI:** 10.1101/2020.06.03.131573

**Authors:** Boris Sotomayor-Gómez, Francesco P. Battaglia, Martin Vinck

## Abstract

Neural coding and memory formation depend on temporal spiking sequences that span high-dimensional neural ensembles. The unsupervised discovery and characterization of these spiking sequences requires a suitable dissimilarity measure to spiking patterns, which can then be used for clustering and decoding. Here, we present a new dissimilarity measure based on optimal transport theory called SpikeShip, which compares multineuron spiking patterns based on all the relative spike-timing relationships among neurons. SpikeShip computes the optimal transport cost to make all the relative spike-timing relationships (across neurons) identical between two spiking patterns. We show that this transport cost can be decomposed into a temporal rigid translation term, which captures global latency shifts, and a vector of neuron-specific transport flows, which reflect inter-neuronal spike timing differences. SpikeShip can be effectively computed for high-dimensional neuronal ensembles, has a low (linear) computational cost that has the same order as the spike count, and is sensitive to higher-order correlations. Furthermore SpikeShip is binless, can handle any form of spike time distributions, is not affected by firing rate fluctuations, can detect patterns with a low signal-to-noise ratio, and can be effectively combined with a sliding window approach. We compare the advantages and differences between SpikeShip and other measures like SPIKE and Victor-Púrpura distance. We applied SpikeShip to large-scale Neuropixel recordings during spontaneous activity and visual encoding. We show that high-dimensional spiking sequences detected via SpikeShip reliably distinguish between different natural images and different behavioral states. These spiking sequences carried complementary information to conventional firing rate codes. SpikeShip opens new avenues for studying neural coding and memory consolidation by rapid and unsupervised detection of temporal spiking patterns in high-dimensional neural ensembles.

## Introduction

Information in the brain is encoded by very high-dimensional “ensembles” of neurons, which encode information with spikes. Populations of neurons can produce specific spike patterns depending on sensory inputs or internal variables [1–8]. With new recording techniques like Neuropixels [9], it has become possible to simultaneously record from thousands of single neurons [10–12]. This offers new opportunities to uncover the relationship between multi-neuron spiking patterns and sensory inputs or motor outputs, yet also poses unique mathematical challenges for the unsupervised discovery of the “dictionary” of neuronal “code-words”.

The notion of information encoding relies on the construction of a distance or dissimilarity measure in an N-dimensional space. For example, the distance between binary strings can be measured using the Hamming distance. In the brain, the distance between two multi-neuron spiking patterns is conventionally based on differences in the firing rates (spike / sec). Using this method, it has been shown for example that high-dimensional neural ensembles span a low-dimensional manifold that relates to a stimulus or behavioral variables in a meaningful way [13, 14]. However, firing rates do not capture the potentially rich information contained by the precise temporal order in which spikes are fired, e.g. neuron *i* firing at time *t* and neuron *j* firing at *t* + *τ*. For instance, we expect that any time-varying sensory stimulus or action sequence may be encoded by a unique multi-neuron temporal pattern of spiking. Indeed, multi-neuron temporal sequences can encode information about sensory stimuli and are required for the generation of complex motor patterns like bird songs [1–7,15,16]. Temporal sequences may also be critical for memory formation, because neural plasticity rules are highly sensitive to the temporal order in which spikes are fired [17–21]. It is plausible that much of the information contained in spiking sequences has thus far not been discovered, as temporal correlations have typically been studied based on relatively small neural ensembles, whereas the number of pairwise spike-time relationships scales with *N* ^2^.

A major computational problem is thus to measure the dissimilarity of spiking patterns in terms of the relative spike timing between neurons. Developing such a measure has several challenges, including 1) Techniques that rely on binning spikes and require exact matches of patterns (e.g. information theoretical measures) have several disadvantages: They require a relatively large number of observations due to combinatorial explosion, lack robustness against spike time jitter and reduce temporal resolution due to binning.

Computational cost becomes a major constraint for high-dimensional ensembles of neurons, and a measure should ideally have a computational cost that is linear in the number of neurons and spikes.

Here, we develop a novel dissimilarity measure for multi-neuron spiking patterns called SpikeShip, which has linear computational complexity of *𝒪* (*N*), and has the key advantage of being sensitive to higher-order structure. SpikeShip can be interpreted as the optimal transport cost to make all spike-timing relationships between two different multi-neuron spiking patterns identical. That is, it solves the optimal transport problem for the entire spiking pattern, and yields a unique decomposition of spike pattern dissimilarity in terms of neuron-specific flows (which controls similarities in terms of relative spike times) as well as a global flow term (which controls the similarity in terms of absolute time). We demonstrated the power of the SpikeShip measure by applying it to large scale, high-dimensional neural ensembles in mice from [12] and [22], and demonstrating that temporal spiking sequences reliably distinguish between natural stimuli and different brain states. We discuss the properties of this measure compared to previous spike train measures like Victor-Purpura Distance (VP) [23, 24], SPIKE [25], and Rate-Independent SPIKE (RI-SPIKE) [26]. Finally, we show that SpikeShip carries orthogonal information compared with the traditional firing rates code.

## Results

𝒪ur overall goal is to develop a dissimilarity measure between multi-neuron spiking patterns that is exclusively based on the temporal order of spiking. Suppose that we have measured the spikes from *N* neurons (“spike trains”), with spike trains divided into *M* epochs of length *T* (measured in seconds or samples). Epochs could be defined by e.g. trials (e.g. stimulus presentations) or sliding windows. The problem is to find a dissimilarity measure with the following properties:

1. The measure should depend on the temporal order of firing across neurons, but not on the spike count.
2. If two spike patterns are identical in terms of cross-neuron spike timing relationships (i.e. they are a temporally translated version of one another), then the dissimilarity measure should equal zero.
3. 3.The measure does not require binning or smoothing and is based on the exact timing of the spikes.
4. It should measure dissimilarity in a gradual way, and avoid the problem of “combinatorial explosion” that occurs with methods that search for exact matches in spiking patterns. Combinatorial explosion means that for a very large number of neurons, the probability of an exact match in spiking patterns becomes extremely small.

We introduce a measure that satisfies these constraints, called the SpikeShip measure (see Methods). The idea of SpikeShip is to measure the dissimilarity between spike trains using the mathematical framework of optimal transport, as shown in Fig. 1 and S1. We will consider each spike train as a collection of “masses” (i.e. the spikes). All spikes from each active neuron, together, contribute a unit mass, i.e. the mass of each spike is normalized to the total mass. This ensures the rate invariance of the method. The question now is what the optimal way is of transporting the masses in time to make the two patterns identical in terms of the relative spike times among neurons. Importantly, similarity here is strictly defined based on *relative* timing among neurons, i.e. not on the absolute timing of the spikes. Intuitively, one would suspect that measuring the similarity of spike train patterns based on the relative spike timing among *N* neurons has a computational complexity of at least order *N* ^2^, which would make the method impractical for larger data sets. However, we surprisingly show that there is a fundamental solution that can be computed in order *N*.

**Fig 1.**
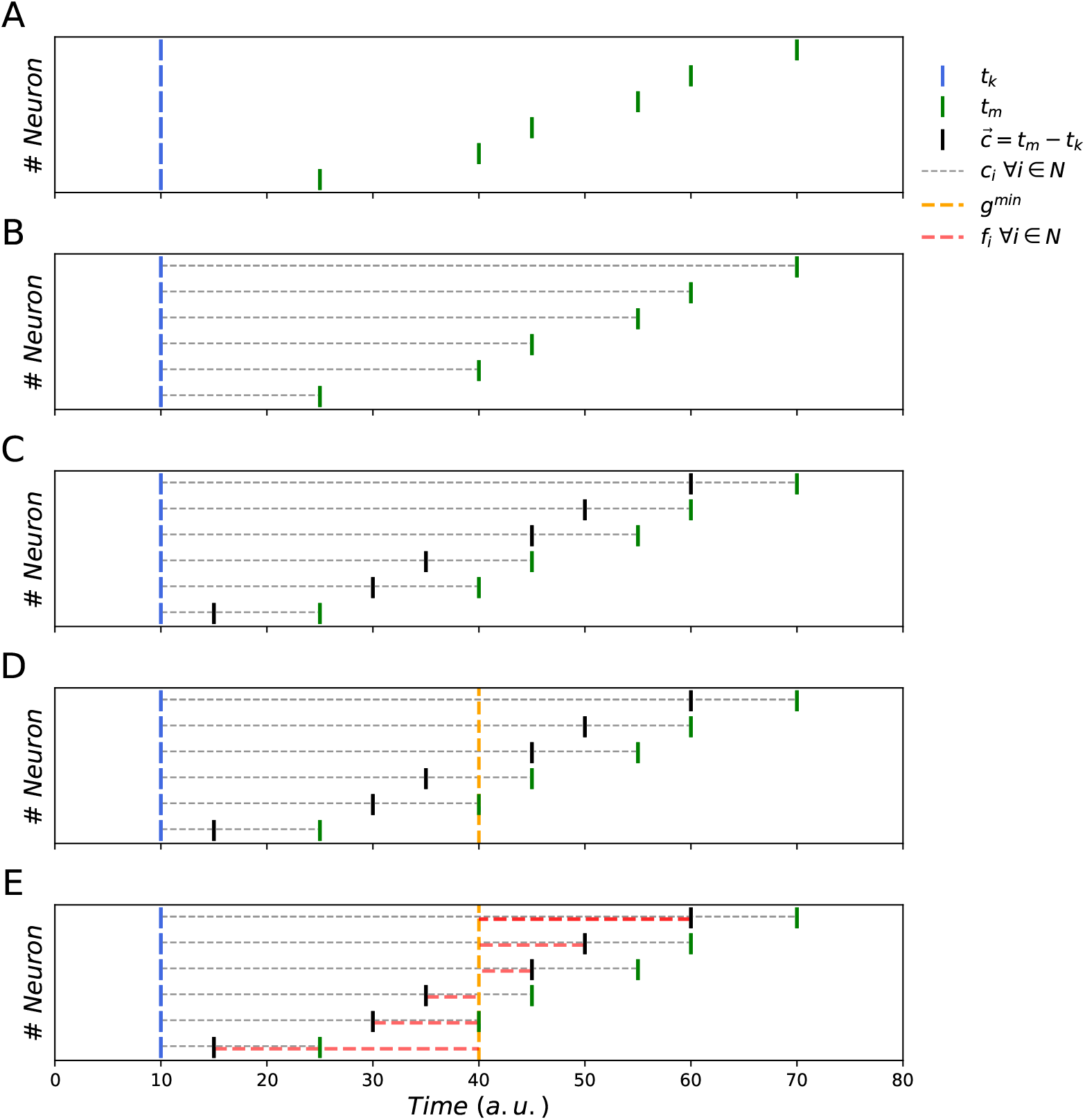
Illustration of SpikeShip. A) Example of two epochs with spike times *t*_*k*_ = (10, 10, 10, 10, 10, 10) and *t*_*m*_ = (25, 40, 45, 55, 60, 70) (note only one spike per neuron in this example). B) Distances between spike times *t*_*k*_ and *t*_*m*_. C) The vector 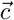 contains the differences of spike times *t*_*k*_ and *t*_*m*_. D) Computation of the median of 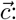 *g*^min^ = 40. E) *g*^min^ is the optimal global shift such that *f*_*i*_=*c*_*i*_ – *g*^*min*^ The neuron-specific shifts 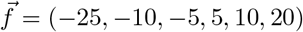 contain all the information about the structure of distances between *t*_*k*_ and *t*_*m*_. SpikeShip equals 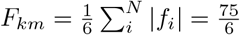

We show that the global optimal transport problem can be solved in two steps:

1. We first compute the optimal transport flow to transform a spiking pattern in epoch *k* into the spiking pattern in epoch *m*, such that the patterns are identical in terms of *absolute* timing.
2. Transport cost is now minimized by computing a global temporal translation term and subtracting this term from all the individual spike shifts. This yields neuron-specific flows, and allows us to compute the total transport cost needed to make two patterns identical in terms of relative spike times.

The algorithm starts by computing the Earth Mover Distance (see Methods, Fig. S1) for each neuron separately in step 1, shifting mass from each spike in pattern *k* to the spikes in pattern *m*. We denote the flows *c*_*i,u*_ with moved mass *w*_*i,u*_, such that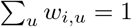

The EMDs, however, only indicate the similarity of absolute spike times between two epochs, i.e. are based on the alignment of the spikes relative to an event. Yet, we wish that our measure reflects the relative timing of spikes between neurons. We show that from the EMD flows computed in Step 1, we can subtract a global rigid translation term. This uniquely yields the minimum transport cost to transform the multi-neuron spiking pattern in epoch *k* such that its relative spike-timing relationships between neurons become identical to another pattern *m*. In the Methods section, we state our main theoretical result, namely that the optimal transport flows are given by

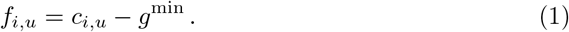

Here *g*^min^ is the weighted median across all the original flows *c*_*i,u*_ with associated mass *w*_*i,u*_. Thus, we can decompose the transport flow in two terms: (1) an optimal global shift between two epochs, shared across all neurons; and (2) an optimal neuron-specific transport flow. We then define SpikeShip (see Methods) as

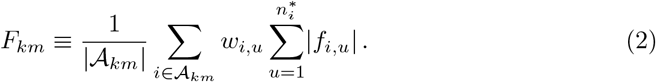

Here 𝒜 _*km*_ is the set of all neurons that fired a spike both in epoch *k* and *m*. The weight 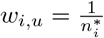 effectively assigns an equal weight to each neuron that contains at least one spike.

The algorithm to compute SpikeShip has computational complexity *𝒪* (*Nn*), because the weighted median has complexity *𝒪* (*N*) (Fig. S2). This means that SpikeShip performs much better in terms of computational complexity than previous measures like SPOTDis (which is *𝒪* (*N*^2^)) [27] and it thus becomes feasible to compute for large ensembles of neurons, as we will show further below.Having computed a dissimilarity measure between multi-neuron spike trains, we can use embedding and clustering techniques to detect patterns in an unsupervised way. The rationale of our approach is that unsupervised clustering can be performed based on the dissimilarity matrices, rather than on the spike train data itself. To illustrate this, we generated 6 input patterns defined by the instantaneous rate of inhomogeneous Poisson processes, as in [27]. Noise was generated with random firing based on a homogenous Poisson process with a constant rate (i.e., homogenous noise) (See Fig. 2A). We computed the pairwise distances between pairs of epochs using SpikeShip distance, yielding a dissimilarity matrix (Fig. 2B). The patterns contained in the data can be visualized using manifold learning algorithms such as t-SNE, using the SpikeShip dissimilarity matrix as input [28–30] (Fig 2C). Furthermore, HDBSCAN [31] automatically detected clusters on the basis of the SpikeShip dissimilarity matrix. These results illustrate how SpikeShip can unveil multi-neuron spiking patterns and shows its efficiency in simulated, high-dimensional data.

**Fig 2.**
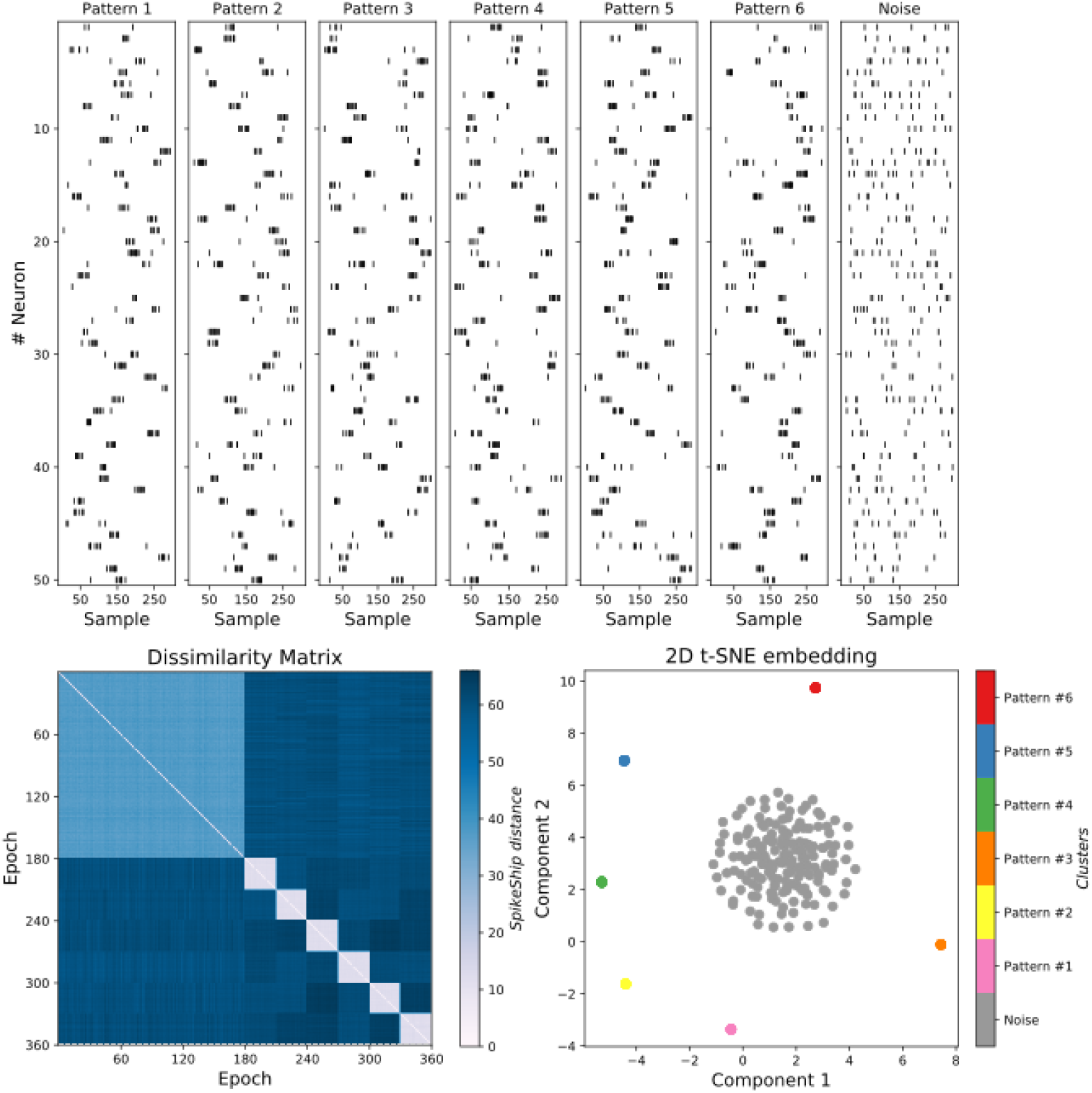
Dissimilarity matrices and clustering comparison. A) Six samples from simulated neuronal outputs according to an inhomogenous Poisson process, plus one for noise (using scripts provided by [27]). B) Top: Sorted Dissimilarity matrix by pattern using SpikeShip dissimilarity measure (left), and a 2-dimensional t-SNE embedding using SpikeShip dissimilarity matrix (right). Bottom: Sorted Dissimilarity matrix by pattern using SPOTDis dissimilarity measure (left), and a 2-dimensional t-SNE embedding using SPOTDis dissimilarity matrix (right).

An important property of SpikeShip is that it can distinguish spiking patterns even when they are multi-modal. To demonstrate this, we use multiple bimodal activation of Poisson patterns with patterned noise and homogeneous noise (See Fig. S3A). In addition, we simulated a special case when neurons are “deactivated” in a certain segment of the epoch (See Fig. S3B). We observed that, in both cases, SpikeShip can detect the patterns successfully and each cluster were well separated as shown in their dissimilarity matrices and 2D t-SNE embeddings (See Fig. S3). Finally, compared to the SPOTDis measure, which is also based on optimal transport (SPOTDis), we found that SpikeShip can detect patterns that are defined by lower signal-to-noise ratios (See Fig. S4).

### Properties of SpikeShip and comparison to other methods

Next, we show that, besides the properties discussed above, SpikeShip has several other desirable features that make it well-suited as a dissimilarity measure for spike train patterns, and that distinguish it from previous spike train metrics:

1. SpikeShip measures the dissimilarity of spiking patterns between a pair of epochs based on the *relative* spike timing between neurons. SpikeShip can therefore be used to detect patterns that are spontaneously occurring or that are not locked to the onset of an event, using e.g. sliding window approaches. By contrast, other measures like VP and SPIKE are not based on the relative spike timing. VP and SPIKE can, however, be used to compare spike trains in different epochs based on the *absolute* spike timing (Figure S5B). In this case, these measures are computed for each neuron separately by directly comparing the spike trains in different epochs, one neuron at a time. To show that SpikeShip can be used in cases where the onset of the pattern was not known, we performed two kinds of simulations. First, we studied a case where the onset of the patterns was not known, and the duration of the pattern was also not known. In this case, a sliding approach could be used to optimize the window length based on Silhouette score. We found that SpikeShip outperforms both SPOTDis and Victor-Púrpura with a sliding window approach, as shown in Figure S6B. Second, we considered a scenario in which there were random global shifts superimposed onto different patterns (See Figure S7A). Here, we added simulations in which there were different patterns, plus global shifts of these patterns relative to the epoch onset. Importantly, the global shifts were not systematically related to the different patterns. As shown in Figure S7B, VP cannot cope with these global shifts, whereas SpikeShip still detects the original patterns based on relative spike timing. In addition, SpikeShip yields the global flow term and thereby directly provides a global measure of latency of the entire pattern.
2. SpikeShip has a linear dependence on differences in relative spike-timing between epochs (Figure S5B). Conversely, because VP has a hyperparameter (*q*), it can have a highly non-linear dependence on spike timing. This is because as differences in timing become larger, VP is exclusively driven by insertions, such that the total cost does not increase (Figure S5B). We further observed that the two other measures, SPIKE and RI-SPIKE, were also influenced by the distance of spiking patterns to the edge of the window (Figure S5B).
3. By design, SpikeShip is only sensitive to timing relationships, and not to firing rates. The reason is that all the spike trains due to normalization have the same mass. By contrast, VP should have a strong rate sensitivity because it also includes the insertion cost (See Fig. S8). In fact, VP will be only rate sensitive when its hyperparameter *q* is very small or very large, and exhibits temporal sensitivity only for intermediate values of *q*. While SPIKE may have some rate sensitivity, RI-SPIKE is designed to have low rate sensitivity [26].

To illustrate these differences, we first examined an example previously shown in [32], with three patterns that either differ by firing rate (1 and 2) or by timing (1 and 2 vs. 3) (Figure S5A). In this example, the VP distance is primarily dominated by rate differences, irrespective of the choice of *q*. In the example shown in Figure S5A, three patterns are shown that either differ by firing rate (1 and 2) or by timing (1 and 2 vs. 3). For all *q*, VP does not assign the lowest between patterns 1 and 2 despite these two patterns having a very close temporal relationship. By contrast, SpikeShip, SPIKE, and RI-SPIKE do assign a much lower cost between patterns 1 and 2 (See Fig. S5A). Additionally, we noted that several measures require temporal alignment. To demonstrate this point, we simulated three Poisson patterns *A, B*, and *C*, as shown in Fig. S5B. Here, the pattern *A* contains Poisson spikes in a specific interval of the window length. Also, we generated the same amount of patterns but after applying a linear shift for half of the neural population and the entire neural population (pattern *B* and *C*, respectively) as illustrated in Fig. S5B. We found that SPIKE and RI-SPIKE are affected by the position of the stimulus onset. Additionally, VP assigns a maximum distance when the cost of shifting spikes is greater than the cost to insert them. Thus, these measures require a clear definition of the stimulus onset to cover due to the definition of the window length affects their distances.

Second, we studied different spiking sequences together with a local or global scaling of firing rates across epochs. As shown in Fig. S8, VP distance did not uniquely distinguish the patterns based on temporal or rate structure, and is dominated by differences in firing rates for a large range of *q*. By contrast, the performance of SpikeShip was not affected by the scaling of firing rates between epochs (See Fig. S9 and S10). Finally, both SPIKE and RI-SPIKE were less effective than SpikeShip as they assign very high dissimilarities between the noise spike trains, such that the overall clustering is worse than for SpikeShip (See Figure S11).

### Application to Allen Brain Institute’s neural datasets

Next, we applied SpikeShip to several high-dimensional neural datasets. Previous work has shown that the visual system can very rapidly process natural images and extract categorical information [33]. It has been proposed that this relies on a temporal coding strategy, whereby visual information is encoded based on the temporal sequence of spikes relative to stimulus onset [33].

Here, we used SpikeShip to determine whether temporal spiking sequences in six visual areas can reliably distinguish natural images from each other. To this end, we analysed Neuropixel data from 32 mice while they passively viewed natural images (dataset from the Allen Institute for Brain Science; see Methods). A total of 20 natural scenes were selected with 10 repetitions each (i.e. *M* = 200 epochs) as shown in Fig. 3A. To create a high-dimensional vector of neurons, we pooled together all recorded neurons (*N* = 8, 301) across the 32 mice (See Fig. 3B).

**Fig 3.**
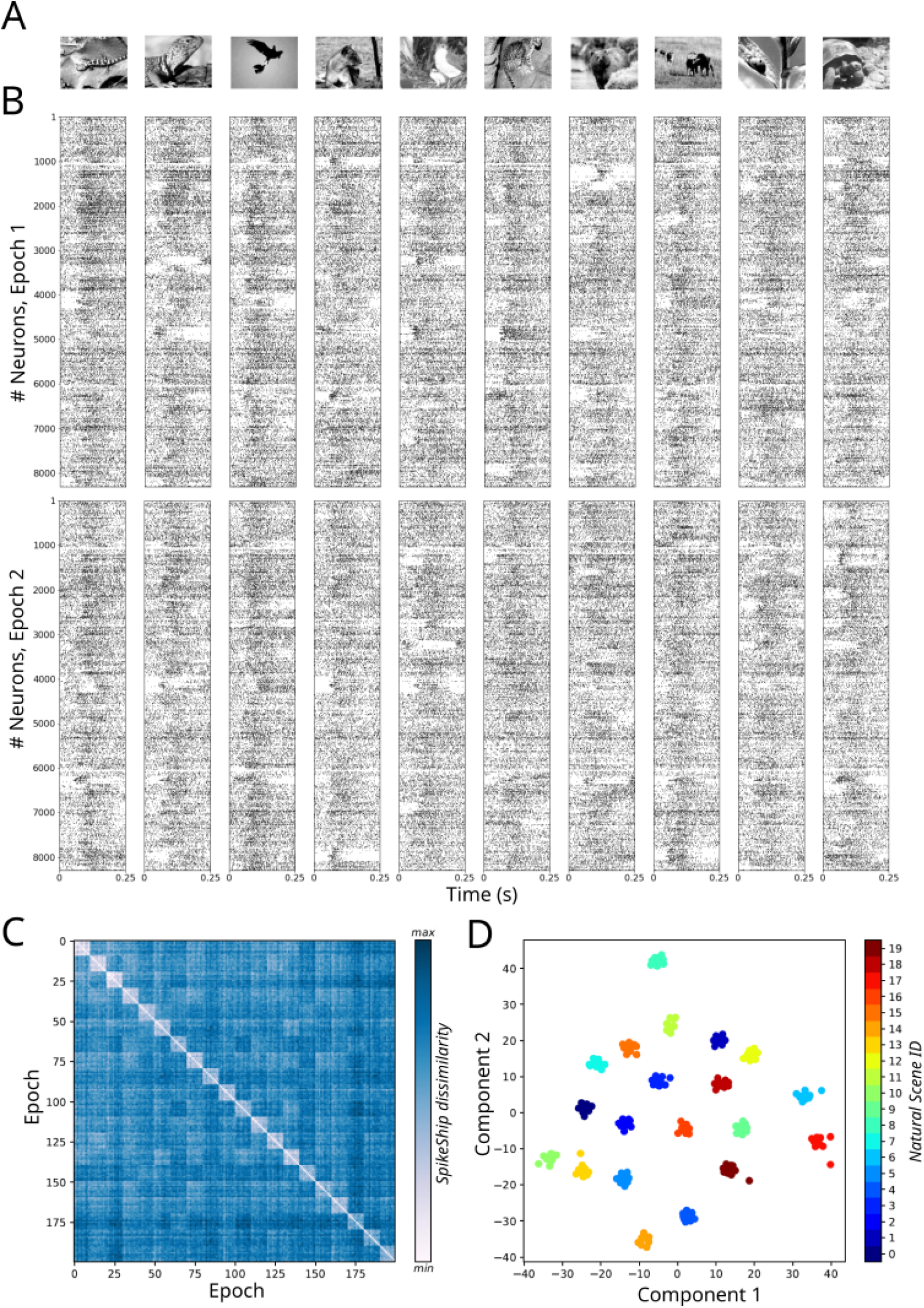
SpikeShip analysis of spike sequences for natural scenes presentations (Allen Brain Institute). A) Sample of 10 Natural Scenes from Allen Brain Institute’s dataset. B) Raster plot of two epochs with *N* = 8, 301 neurons for each presentation. C) Left: Sorted dissimilarity matrix by image ID for 20 natural scenes presentations with 10 presentations each (*M* = 200). Right: 2-dimensional t-SNE embedding for each presentation.

The pairwise SpikeShip distance is presented in Fig. 3C as a dissimilarity matrix sorted by the presentations of each natural scene. The t-SNE embedding revealed clear clustering of spiking patterns based on SpikeShip, such that different natural images could be reliably distinguished from each other (See Fig. 3D). Hence, natural images yielded distinct temporal spiking sequences that were time-locked to stimulus onset (so that they could be extracted from combined data from multiple sessions), in support of the idea that the visual system may use a temporal coding strategy. In sum, these findings demonstrate that SpikeShip can unveil multi-neuron temporal spiking patterns from high-dimensional recordings.

### Comparison of SpikeShip vs firing rates in visual stimuli

We wondered how the information in temporal spiking sequences compared to the information carried by conventional firing rate codes. Our first question was, which one of these two codes conveyed more information. To determine this, we computed a distance matrix for the firing rates, by computing the Euclidean distance between the firing rate vectors. These distances were computed in the same time window as we used for SpikeShip (Fig. 4A). We then computed a measure of discriminability between natural images based on the distances within and between images (see Methods, Eq. 26).

**Fig 4.**
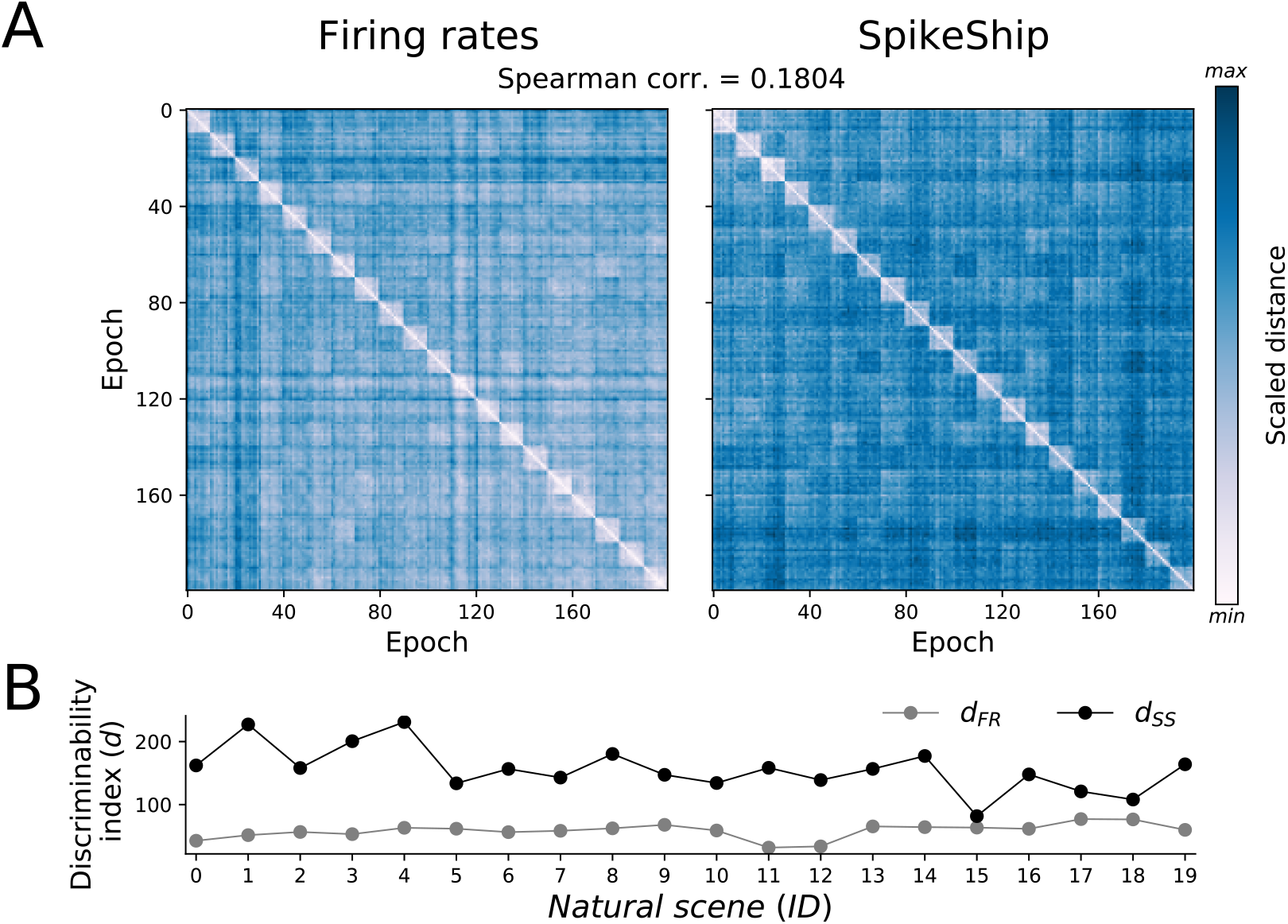
Comparison of pairwise distances between Firing rates and SpikeShip. Dissimilarity matrices for Allen Brain Institute data (sorted by natural scenes). Colormaps were changed to show more variability in distances. B) Discriminability index for dissimilarity matrices shown in A) by using firing rates (*d*_*F R*_) and SpikeShip (*d*_*SS*_).

We found that there was a higher discriminability between natural images for SpikeShip as compared to firing rate vectors for all the natural scenes (Fig. 4B).

Next, we wondered to what extent the information in SpikeShip was independent of the firing rate information. To this end, we computed the Spearman correlation between the SpikeShip distance and firing rate dissimilarity matrices, which contained information from all epochs across all natural images. We found that the dissimilarity matrices of SpikeShip and firing rates were only weakly correlated across epochs (Spearman correlation equals 0.1804). Accordingly, the t-SNE visualization shows that the relative locations of the clusters show major differences between both methods (Fig. S12).

Altogether, these results show that SpikeShip and firing rates contain different and to a large extent independent information about natural stimuli. Furthermore, SpikeShip allowed for a better separation of the different natural stimuli in comparison to firing rates. We further observed that both SPIKE and RI-SPIKE were less effective in separating the different patterns than SpikeShip, and showed a stronger correlation with firing rates (Fig. S13). These findings support the idea that the visual system may use spiking sequences to encode information about natural scenes, as proposed by e.g. [33].

### Application to spontaneous activity

Next, we applied SpikeShip to high-dimensional neural recordings from multiple brain areas while mice spontaneously transitioned between different behavioral states. Previous works have shown that behavioral states have major effects on the firing rates of neurons across multiple brain areas [10, 34]. Recent work has shown that different facial motion components outperform the prediction of firing rates as compared to running speed and pupil diameter (a measure of arousal). Hence, we wondered whether different behavioral states are accompanied and distinguished by specific spiking sequences.

To investigate this, we analysed the data set of [22], which contains multi-areal recordings from *>* 1000 neurons in three mice. Similar to [10], we distinguished between different states based on the facial motion components using the SVD (singular value decomposition), and identified low, medium, and high-motion epochs (Figure 5A-C). We randomly selected one hundred epochs for both low-, medium and high-motion states and computed the SpikeShip dissimilarity matrix (*M* = 200, See 5D).

**Fig 5.**
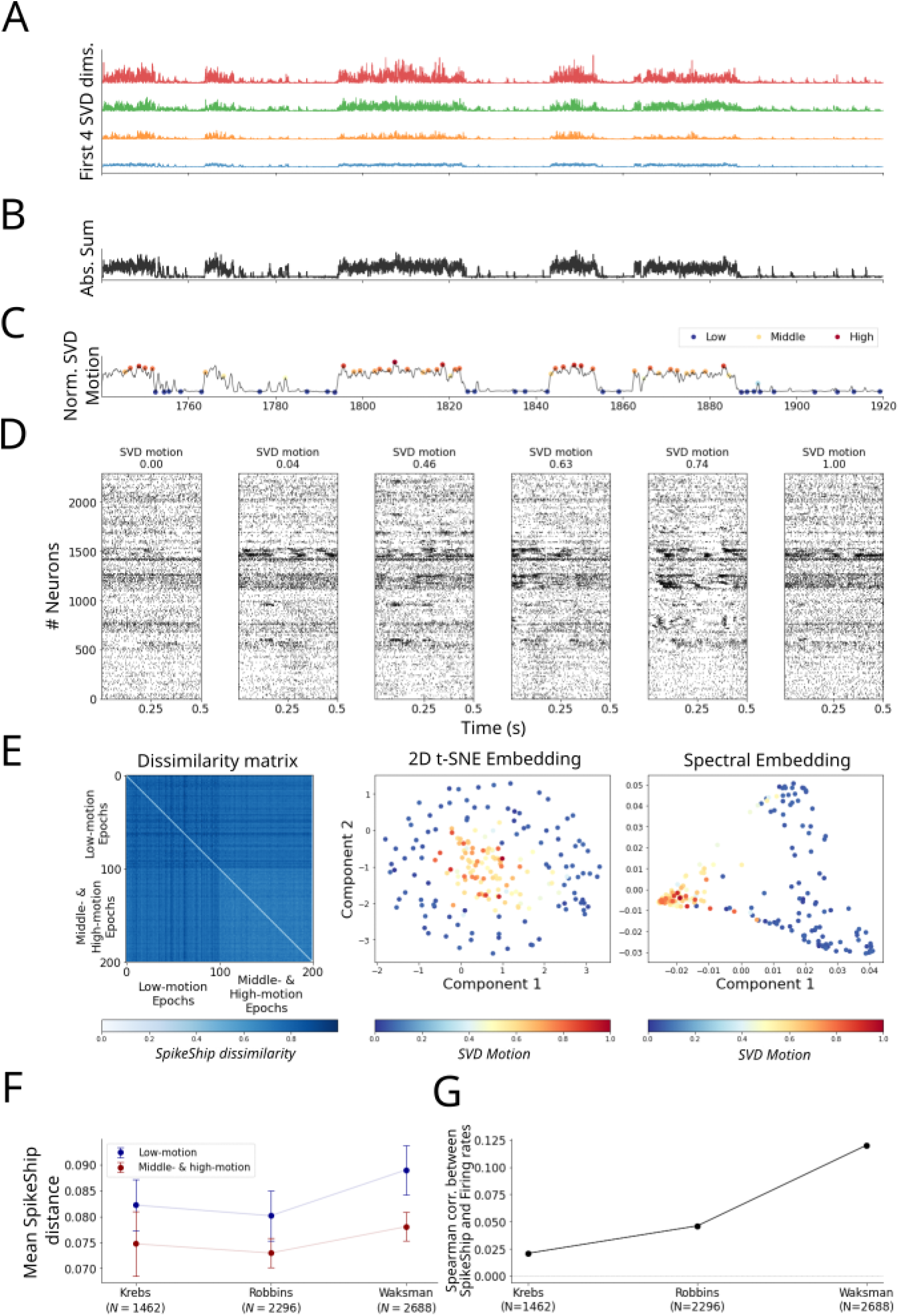
Spontaneous activity analyses. A) First 4 SVD dimensions of [22] dataset. subtraction of the median of the first four SVD motion dimensions (denoising). C) Normalized SVD motion used to detect low- and high-states. Circles represent samples of temporal windows for posterior analyses. D) Raster plot for *N* = 2, 296 neurons for different epochs. E) Multi-spike sequence analyses. Left: Dissimilarity Matrices. Middle: 2D t-SNE embedding. Right: 2D Spectral Embedding (Laplacian Eigenmaps). F) Mean SpikeShip distance from [10] experiments. G) Spearman correlation between firing rates and SpikeShip per experiment.

In Fig. 5E, we show the dissimilarity matrix for one mouse (“Waksman”; *N* = 2, 688, for the other two mice see Fig. S14). The first 100 epochs represent low-motion states, and the remaining 100 are the middle- and high-motion epochs. The dissimilarity matrix reveals that the spiking sequences during medium and high motion are relatively similar to each other, whereas there is a relatively high variability among sequences during low-motion epochs (Fig. 5E-F). Furthermore, both the t-SNE embedding and the spectral embedding show a separate state-space region for the spiking sequences during medium and high motion.

We further wondered if SpikeShip contains orthogonal information compared to firing rates, similar to what we had observed for natural images. Again, we computed the Spearman correlations between the SpikeShip and the firing rate dissimilarity matrices. We found relatively weak correlations between the firing rate and SpikeShip dissimilarity matrices for the three mice: (0.189, 0.024, *−* 0.064) for *N* = (1462, 2296, 2688) neurons, respectively (See Fig. 5G).

Altogether, these findings indicate that different brain states give rise to specific temporal correlation patterns across neurons, with relatively homogeneous spiking sequences during active behavior as compared to quiescence.

## Discussion

We studied the problem of measuring the dissimilarity between two multi-neuron spiking patterns based on the relative spike-timing relationships across neurons (i.e. firing order). We developed a new measure called *SpikeShip*. SpikeShip solves the problem of optimally transporting the spikes of individual neurons, such that the global pattern of spike-timing relationships becomes identical between two epochs. Intuitively, one would think that such a measure has a computational complexity of at least *𝒪* (*N* ^2^), but we show that it can be computed with *𝒪* (*N*) computational complexity; this is a major improvement over our previous work [27]. We show that a dissimilarity between two spiking patterns can be decomposed into neuron-specific flows and a temporal rigid translation term. Importantly, SpikeShip is not restricted to the 2nd order correlations, but is based on the higher-order structure in the spike train.𝒪ur technique can also be used to align temporal sequences in an unsupervised way if there is a global jitter between spike patterns. This achieves a similar goal as a recent study [35], described as “warping”; however, the method in our paper does not require any parameter estimation; furthermore, it can also be used to align trials corresponding to different stimuli or behavioral conditions. For example, suppose that we want to analyse the sequence of neural activation in relationship to some event, e.g. freely moving your arm. Suppose that we do not know the exact onset of the neuronal sequence in relation to moving your arm, and we record multiple realizations of this event. With SpikeShip, we can immediately decompose the transport cost between any two trials in terms of a global translation term (which is a non-linear computation) and a neuron-specific shift. We showed here that the global translation term picks up shared latency shifts between neurons. The global translation term allows one to align trials that were recorded in different conditions, e.g. moving your arm left or right. And it yields, in essence, a state-space characterization of temporal trajectories (e.g. related to different movements), rather than a trajectory through a state-space of firing rate vectors.

We applied SpikeShip to large, real neuronal datasets of experiments in mice. We found that SpikeShip can be used for the unsupervised decoding of different natural images from a high-dimensional temporal spiking pattern across *N >* 8000 neurons. Interestingly, we found that SpikeShip outperformed the classical firing rate vector, and that the spike timing information was only weakly redundant with the information in the firing rate vector. This suggests that spike timing information carries additional information relative to the firing rate, as has been hypothesized by [33]. Interestingly, the SpikeShip technique does not require the explicit identifications of spike latencies, and is able to extract higher-order correlations from spiking patterns and also distinguish multimodal patterns from each other [27]. Furthermore, as we showed here, the computation is extremely efficient. We further analysed large-scale recordings from the visual cortex, retrosplenial, sensorimotor, frontal, striatum, hippocampus, thalamus, and midbrain. We showed that the temporal structure of spike trains distinguished between low- and high-motion epochs, and again provided orthogonal information to the firing rate code. Furthermore, we found that spiking patterns become more homogeneous during high-motion epochs, which is consistent with the notion that arousal improves the reliability of signal transmission in the cortex [34].

SpikeShip has several advantages or distinct properties as compared to other measures: 1) It is useful to note that there are different measures that are designed for distinct computational problems. SpikeShip is explicitly designed to measure the similarity between patterns in terms of relative spike time relationships. This distinguishes it from e.g. the VP distance, which measures similarity in terms of absolute spike times. We note that while SpikeShip is designed to measure similarity based on relative spike timing, it can also be used to measure similarity in terms of the absolute timing of spikes. In doing so, it has the advantage of extracting separately a global translation term, indicating shared latency shifts, and inter-neuronal timing differences. We furthermore showed that SpikeShip can be used in combination with a sliding window approach. Here the computational cost offers a great advantage as many different window lengths can be compared based on e.g. Silhouette score, as we show here. 2) SpikeShip is by design not sensitive to global or local scaling of firing rates, as opposed to VP. We note that also RI-SPIKE is designed to be insensitive to a scaling of firing rates. We note however that SpikeShip dissimilarity between any two epochs is based on the spike trains of those neurons that fire at least one spike. Thus, although the value of SpikeShip is not biased by firing rate, the firing rate can influence which neurons the measure is computed over, in particular when neurons have very low (baseline) firing rates. 3) SpikeShip has the major advantage of finding sequences based on higher-order temporal structure based on relative spike timing, with a computational cost of (*𝒪Nn*). 4) As shown here, SpikeShip is highly noise robust, and outperforms our previous method SPOTDis. 5) SpikeShip can be used for the detection of multi-modal patterns, which e.g. distinguishes it from methods that detect sequences based on the latency of firing. 6) SpikeShip is a bin-less measure, i.e. it is based on exact spike timings. 7) SpikeShip does not require exact matches between patterns, but is based on a metric distance (earth mover distance) that reflects the magnitude of differences in relative spike timings. This is an important difference relative to measures that e.g. require exact matches in patterns. Yet it also distinguishes it from e.g. VP, which shows a highly non-linear dependence on timing differences, and SPIKE and RI-SPIKE, which show a non-monotonic dependence on timing differences.We note that SpikeShip has an efficient computation time of order *𝒪Nn* (number of neurons times number of spikes), which is comparable to the computational cost of the spike count. This is remarkable given that SpikeShip quantifies the dissimilarity based on all the relative spike-time relations. SpikeShip achieves this computation time by computing the Earth Mover Distance (EMD) first for each spike train separately, and obtaining individual flows by computing a global flow. We note that EMD was also applied to cross-correlations (in [27]) and to individual spike trains [36]. In [36] the EMD is quantified one neuron at a time, i.e. without considering inter-neuronal spike-time relationships, and the dissimilarity of spiking pattern is thus based on the absolute timing relative to a stimulus onset. Crucially, SpikeShip aims to quantify the dissimilarity of spiking patterns (between two epochs) in terms of the spike-timing relationships among *all* recorded neurons (i.e. inter-neuronal spike time relationships). This means that SpikeShip is based on relative spike-time relationships, which makes it invariant to e.g. the onset of a sequence relative to the beginning of an epoch, and allows for the quantification of spontaneously occurring sequences that are not locked to a stimulus onset. Thus, SpikeShip allows for a wide variety of applications (including sequences time-locked to a stimulus onset) and is thus more generic than EMD computed per neuron separately.

Looking forward, SpikeShip opens new avenues to study temporal sequences in high-dimensional neural ensembles. Recent technological developments now allow for recordings of thousands of neurons simultaneously, either using electrical recordings or two-photon imaging [9]. The technique developed here is applicable to both kinds of data, due to linear computation time. Application of SpikeShip to such kind of data might generate important insights into the role of temporal sequences in sensory coding and memory consolidation.

## Materials and methods

### Derivation of SpikeShip

Here we derive a new dissimilarity measure, called SpikeShip, which has computational complexity *𝒪* (*N*) for one pair of epochs. To derive this measure, we first consider the simplified case where each *i*th neuron fires one spike for all *M* epochs. In SpikeShip, we use an L1 cost on differences in spike timing, which has two principal reasons: First, using the L1 norm indeed allows for an efficient computation of SpikeShip with computational complexity *𝒪* (*nN*) (number of spikes times number of neurons). Second, using the L1 norm instead of the L2 norm avoids over-weighing large spike time shifts (i.e. a shift from e.g. 0 to 0.1 is weighted similarly as a shift from 1 to 1.1).

### SpikeShip for a single spike per epoch

Let 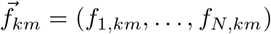be a vector of flows for each neuron in epoch *m*. In other words, *f*_1,*km*_ is the shift of the spike fired by the first neuron in the *m*th epoch. The total moving cost equals

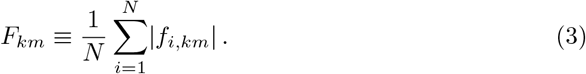

The problem statement is to find a flow vector 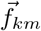 such that after moving the spikes, the resulting spike train patterns are identical between epoch *k* and epoch *m*, in terms of the full matrix of spike timing relationships. We wish to find the flow vector that satisfies this constraint with minimum cost *F*_*km*_.

**Example:** Suppose there are two epochs for *N* = 6 neurons with spike times 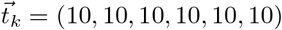 and 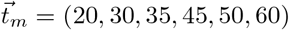. We will show that the flow vector with minimum cost, such that spike patterns have identical temporal structure, equals 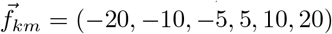 (Fig. 1).

More formally, let *t*_*m,i*_ be the timing of the spike for the *i*th neuron in the *m*th epoch. We denote the spike times after (post) moving them in epoch *m* as

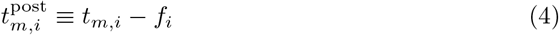

*∀i*, where we omitted the subscripts *k, m* from the variable *f*_*i,km*_ for simplicity. The constraint that all the across-neuron spike timing relationships should be identical after moving implies that*m,i*

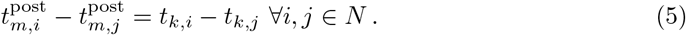

In other words, the delay between two spikes from two different neurons (*i, j*) should be identical between pattern *k* and *m* after moving the spikes. Substituting based on Eq. 4, this can be expressed as

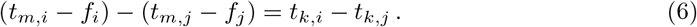

Let *c*_*i*_ be the shift in spike timing for each neuron in epoch *m*, such that the spike train patterns become identical,

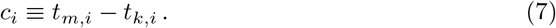

Note that with this definition of *c*_*i*_, the equation

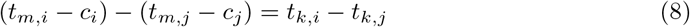

holds for all (*i, j*). For all *i*, we can express *f*_*i*_ as a function of the shift *c*_*i*_, such that

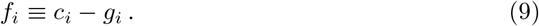

We wish to solve

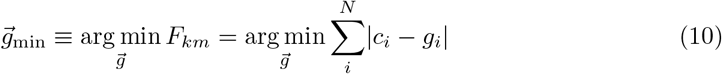

under the constraint of Eq 6.

From Eqs 6, 8 and 9 it follows that

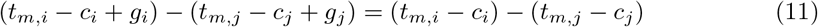

Hence the equation *g*_*i*_ = *g*_*j*_ holds for all (*i, j*). Thus, we can rewrite Eq. 10 to

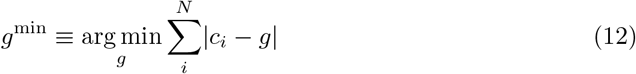

to find a global shift that minimizes the L1-norm of the residuals. The solution to this equation is

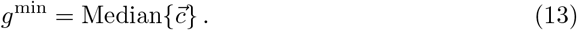

Thus, our main result is that the original shifts between the two spiking patterns can be written as the decomposition

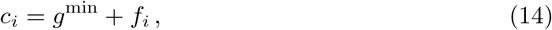

for all *i*, i.e. the optimal transport in terms of neuron-specific shifts and a global temporal rigid translation term. Thus, the optimal transport between two spiking patterns (e.g. after stimulus onset) can be decomposed into the optimal transport in terms of neuron-specific shifts and a global temporal rigid translation term.

### Global shift definition for multiple spikes

We now consider the case where in each epoch, every neuron fires a variable number of spikes. We will show that a similar derivation for SpikeShip can be made based on the weighted median. Let *n*_*i,k*_ and *n*_*i,m*_ be the number of spikes for the *i*-th neuron in epoch *k* and *m*. We will also assign a weight to each spike, such that the total weight per neuron in the computation of SpikeShip is equal. In order to do so, we first find the smallest common multiple of the spike counts for each neuron, denoted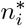. For instance, if *n*_*i,k*_ = 2 and *n*_*i,m*_ = 6 then the least common multiple equals 6. We now replicate each spike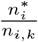 and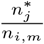 times. Each spike will now have a weight of 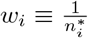. (Note that in the actual computation, we do not replicate the spikes in practice, but use an algorithm similar to the one detailed in [27]). Then, based on the optimal transport cost (EMD), we obtain shifts *c*_*i,u*_ for the *u*-th spike of the *i*-th neuron, 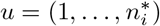. Note that we have shown EMD for spike trains can be efficiently computed by first shifting the mass from the most left-ward spike (i.e. first spike) out of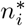 spikes in epoch *m* to the most left-ward spike in epoch *k* [27], and then proceeding with the second spike, etc.

We use a similar derivation as the one above. Let *t*_*m,i,u*_ be the timing of the *u – th* replicated spike for neuron *i* in the *m*th epoch, for all (*i, m, u*), 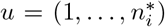. We denote the spike times after (post) moving them in epoch *m* as

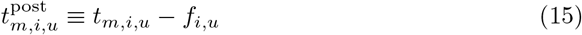

*∀*(*i, u*), where we omitted the subscripts *k, m* from the variable *f*_*i,km,u*_ for simplicity. The constraint that all the across-neuron spike timing relationships should be identical after moving implies that

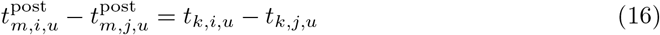

In other words, the delay between two spikes from two different neurons (*i, j*) should be identical between pattern *k* and *m* after moving the spikes. Substituting based on Eq. 15, this can be expressed as

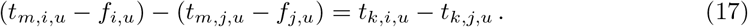

There is one additional constraint, namely

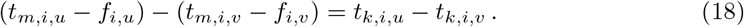

In other words, all the pairwise relationships after moving within the same neuron should be identical.

Let *c*_*i,u*_ be the shift in spike timing for each neuron in epoch *m*, such that the spiking patterns in the window become identical,

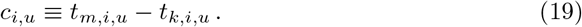

Note that with this definition of *c*_*i,u*_, the equation

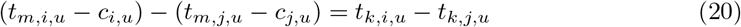

holds for all (*i, j, u*). For all *i, u*, we can express *f*_*i,u*_ as a function of the shift *c*_*i,u*_, such that

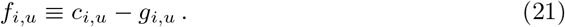

Define 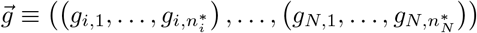

We wish to solve

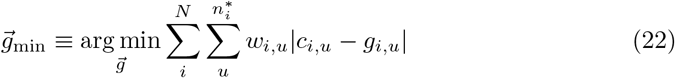

under the constraints of Eq 17, 18 and 20.

Given these two constraints, the equation *g*_*i,u*_ = *g*_*j,u*_ holds for all (*i, j, u*). Thus, we can rewrite Eq. 22 to

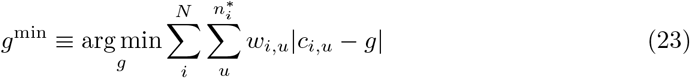

to find a global translation term that minimizes the L1-norm of the residuals. The solution to this equation is the weighted median *g*,

*g*^min^ = WeightedMedian 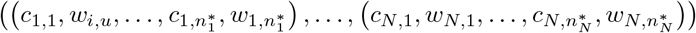

Now average mover cost, *SpikeShip*, equals

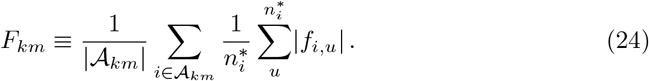

where 𝒜 _*km*_ *i* : *n*_*i,m*_ *>* 0 *n*_*i,k*_ *>* 0 is the set of neurons that are active in both epochs *k* and *m*. Thus, we derive the result that the original shifts between the two spiking patterns can be written as the decomposition

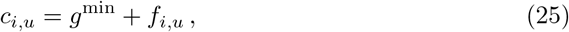

for all (*i, u*), i.e. the optimal transport in terms of neuron-specific shifts and a global translation term. Thus, the optimal transport between two spiking patterns (e.g. after stimulus onset) can be decomposed into the optimal transport in terms of neuron-specific shifts and a global temporal rigid translation term. This computation has linear complexity *𝒪*(*N*). We base the computation of the weighted median on [37, 38] and an adaptation of the *Robustats* [39] Python library. In practice, the computation of *c*_*i,u*_ (optimal transport per neuron) can be efficiently performed not by replicating the spikes to the common multiple integer (used in the derivation above), but by using a similar algorithm as in [27].

### Comparison with other spike train metrics

#### Victor-Púrpura distance

Victor-Púrpura metric (VP) combines both rate and temporal information by defining a hyper-parameter *q* related with the cost of shift between spikes. Thus, VP extracts rate information for small values of *q*, converging to the absolute difference of spike counts for two spike trains with *n*_*i*_ and *n*_*j*_ spikes when *q* = 0 (i.e., *V P* (*q* = 0) = | *n*_*i*_ *−n*_*j*_|). 𝒪n the other hand, for high values of *q*, VP distance maximizes the contribution of timing coding, converging to the sum of the total spike of both spike trains (i.e., *V P* (*q →∞*) = *n*_*i*_ +*n*_*j*_)).We used *Elephant* (Electrophysiology Analysis Toolkit) [40] to compute VP distance.

#### SPIKE and RI-SPIKE distances

Both SPIKE and RI-SPIKE measure the similarity of two spike trains in terms of the absolute spike timing. RI-SPIKE was developed to avoid a rate bias in the computation of the dissimilarity. To perform the analyses for SPIKE and RI-SPIKE metrics, we used a python library for spike train similarity analysis called *PySpike* [41].

#### SPOTDis

Previously, we have developed a dissimilarity measure between multi-neuron temporal spiking patterns called SPOTDis (Spike Pattern Optimal Transport Dissimilarity) [27]. SPOTDis is defined as the minimum energy (optimal transport) that is needed to transform all pairwise cross-correlations of one epoch *k* into the pairwise cross-correlations of another epoch *m*. This optimal transport is given by EMD (Earth Mover Distance). SPOTDis only measures pairwise correlations and has computational cost of order *N* ^2^. We used SPOTDis python module [27] to perform the analyses.

### Application in high-dimensional neural data

#### Allen Brain Institute datasets

We used the free, publicly available datasets of Allen Brain Institute through AllenSDK (For more details, see http://help.brain-map.org/display/observatory/Documentation). Neuropixels silicon probes [9] were used to record neurons with precise spatial and temporal resolution [12].

We selected the cells of 32 mice during natural scene presentations. The cells were selected considering a signal-noise ratio (*SNR*) such that *SNR >* 0. The neural activity from a total of *N* = 8, 301 cells was selected from the Primary visual area (VISp), Lateral visual area (VISl), Anterolateral visual area (VISal), Posteromedial visual area (VISpm), Rostrolateral visual area (VISrl), and Anteromedial visual area (VISam).

The computation time for SpikeShip to compute one dissimilarity matrix of Fig. 3C was on average 58.8 secs (see Methods), thus indicating a highly effective computation time for *N* = 8, 301 neurons. In contrast, we estimate the computation time for the previous SPOTDis measure [27] to be approximately 175.71 hours (1 week and 7.7 hours; based on *n ≈* 3.33 spikes per neuron).

#### Spontaneous activity

For the spontaneous activity dataset, we used free, publicly available datasets [22]. It contains data about three mice: “Waksman” (*N* = 2, 688), “Robbins”(*N* = 2, 296), and “Krebs” (*N* = 1, 462). Probes were located in distinct cortical areas (visual, sensorimotor, frontal, and retrosplenial), hippocampus, thalamus, striatum, and midbrain [10]. Furthermore, each experiment has information about the recorded spike times, cells, and the processed SVD components of whisker’s motion (i.e. pixel difference between consecutive frames of a mouse face movie).

For our analyses, we added the first four SVD components (Fig. 5A) and then we subtracted the median across dimensions (denoising, see Fig. 5B). A sliding window was used to sum the activity SVD the whisker’s motion across the interval [*t, t* + 0.5] seconds (Fig. 5C,D). We selected one hundred epochs for both low- and high-motion states based on the SVD motion value, and ran SpikeShip across different amounts of cells, depending on the experiment. The manifold learning algorithms were t-SNE and Spectral Embedding from the python library Scikit Learn v0.22.1.

For these experiments, SpikeShip took 25.5 secs in creating a dissimilarity matrix. Considering *N ≈* 2, 500, *n ≈* 5, based on Fig. 5D, the computation time of SPOTDis can be estimated as 34.06 hours (1 day and 10.06 hours; *SU ≈* 4, 808).

#### Discriminability index

Since the information in temporal spiking sequences compared to the information from traditional firing rate codes could differ, we wanted to quantify to what extent the degree of discriminability within and between different visual stimuli differ in a dissimilarity matrix *diss* (i.e. within and between natural images) per stimulus *id*.

To this end, we computed a “Discriminability index” (*d*), defined as the difference between the average distance within and between stimulus *id*, divided by the squared sum of their variances to the power of two.

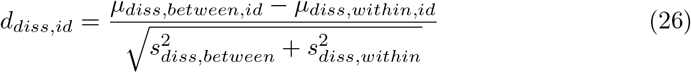

Therefore, it indicates how many standard deviations are two sets of distances away from each other.

## Code availability

The source code of *SpikeShip* can be accessed from the GitHub repository: https://github.com/bsotomayorg/SpikeShip.

## Acknowledgements

This project was financed by the BMF (Bundesministerium fuer Bildung und Forschung), Computational Life Sciences, project BINDA (031L0167).

## Authorship Contributions

Conception and problem statement: BSG, FPB, MV. Data analysis and simulations: BSG. Mathematical analysis: BSG and MV. Supervision: MV. Writing of main draft: BSG and MV.

## Supporting information

**Fig S1.**
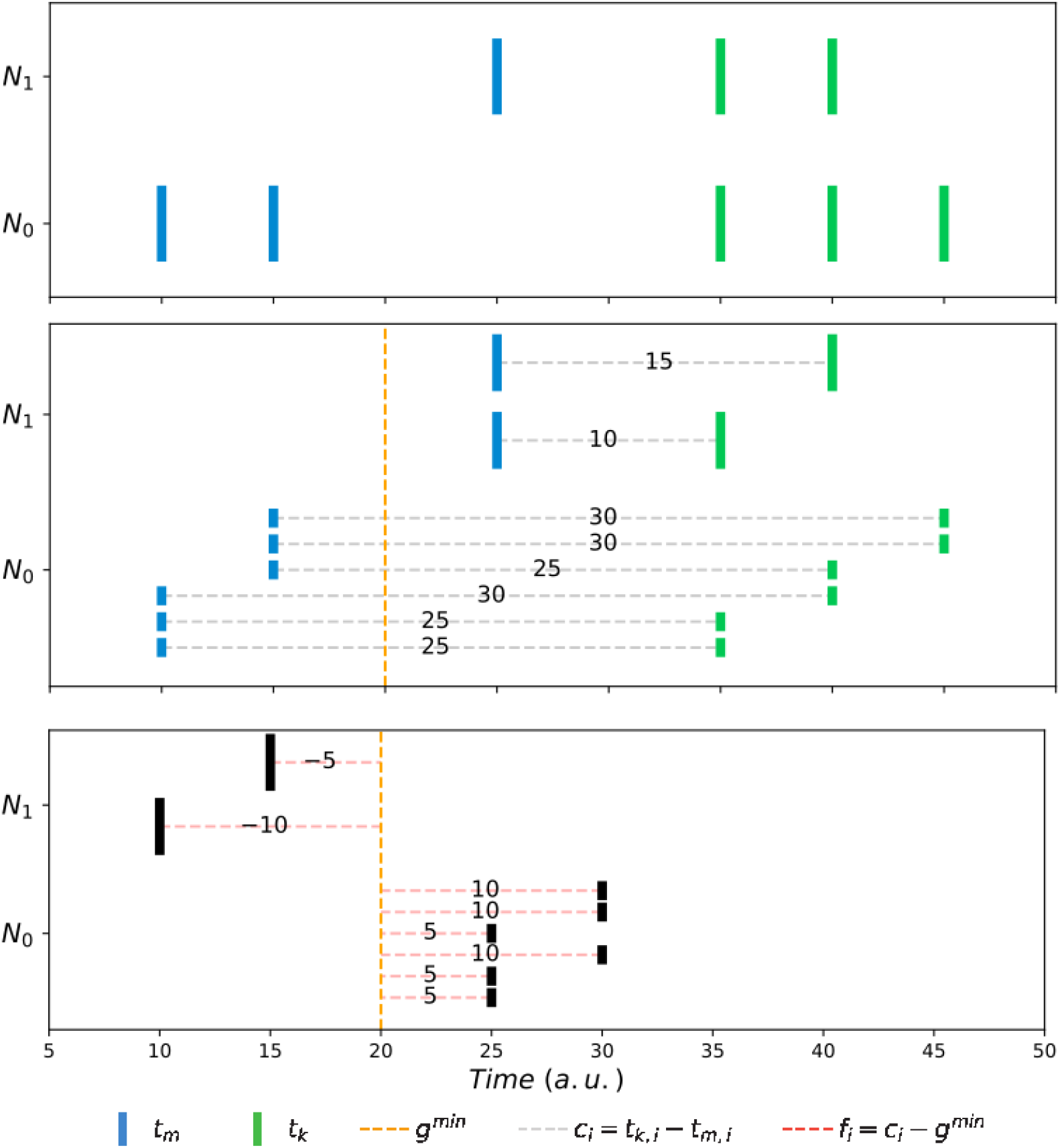
Illustration of SpikeShip for multiple spikes. (Top) Example of two epochs with spike times for two active neurons *N*_0_ and *N*_1_ (i.e., *A*_*km*_ = 2): *t*_*k*_ = ((10, 15), (10)) and *t*_*m*_ = ((35, 40, 45), (35, 40)). Spike counts per epoch *m* and *k* correspond to (*t*_*N*_0 _,*m*_ = 2, *t*_*N*_1 _,*m*_ = 1) and (*t*_*N*_0 _,*k*_ = 3, *t*_*N*_1 _,*k*_ = 2), respectively. (Middle) The difference of spike times is computed by normalizing the mass across neurons and between epochs. Such spike time difference is 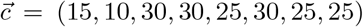with mass (i.e., weights) 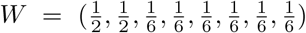. Then, the global shift (i.e., *g*^*min*^) equals 20. (Bottom) Neuron-specific shifts correspond to 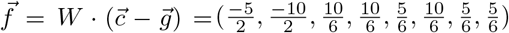. Thus, 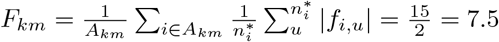

**Fig S2.**
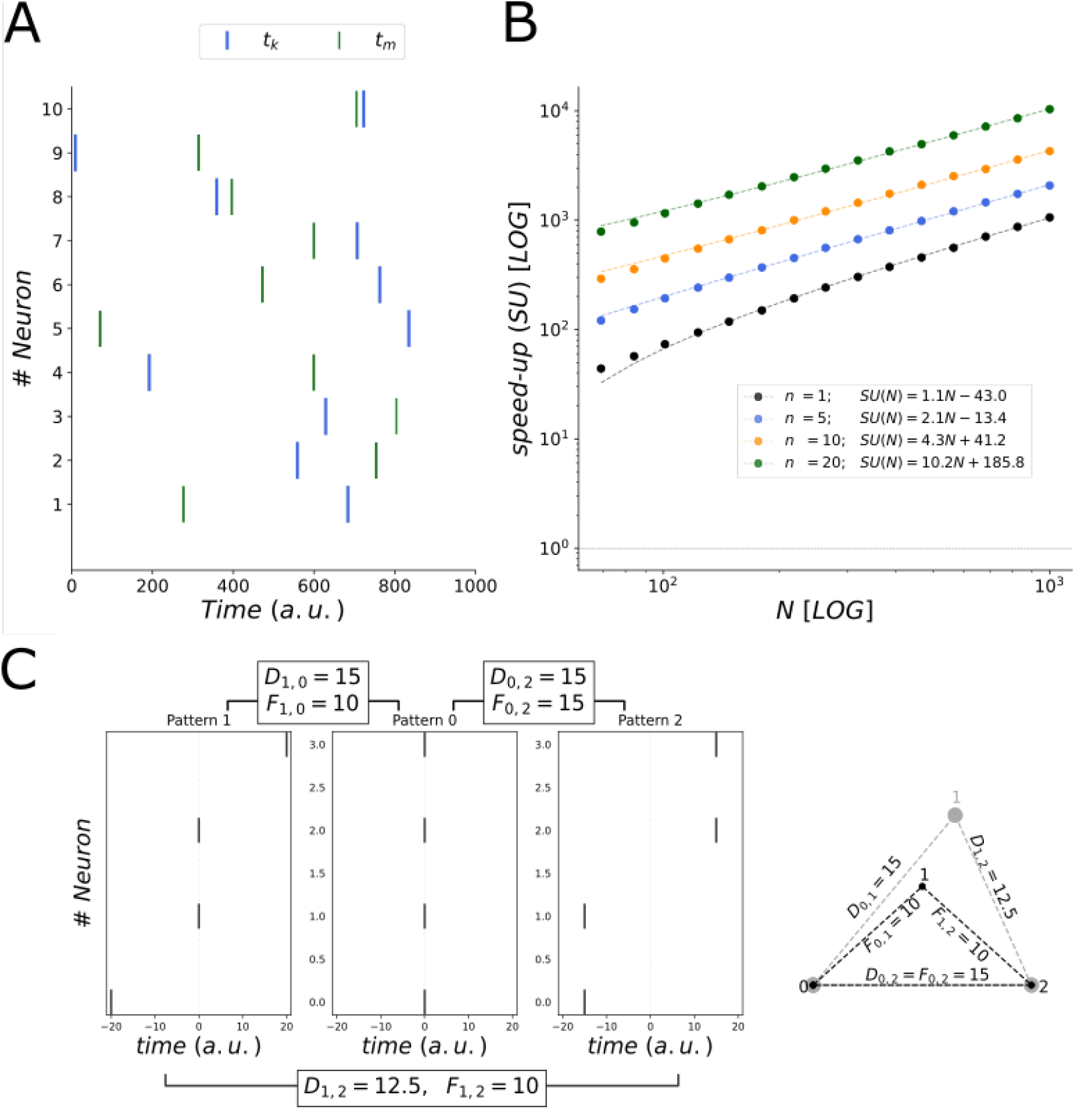
Accuracy and Speed-up comparison for single- and multi-spike patterns. A) Example of single spike trains for two epochs for 10 neurons. Patterns were generated as uniform sequences with *n* = 1 spike per neuron per epoch. B) Computational speed-up (log-scale) for SpikeShip vs. SPOTDis (serial execution) grossberger2018unsupervised for increasing amount of neurons *N*. Speed-up is approximately *N* when there is 1 spike per neuron, and it increases when *n >* 1 (i.e. the multi-spike pattern case). C) Example of three single-spike patterns: (*−*20, 0, 0, +20), (0, 0, 0, 0), and (*−*15, *−*15, +15, +15), from left to right. SpikeShip assigns a geometrically more appropriate transport cost between pattern 1 and 2 (*F*_1,2_ = 10) than SPOTDis (*D*_1,2_ = 12.5), considering their distance with pattern 0.

**Fig S3.**
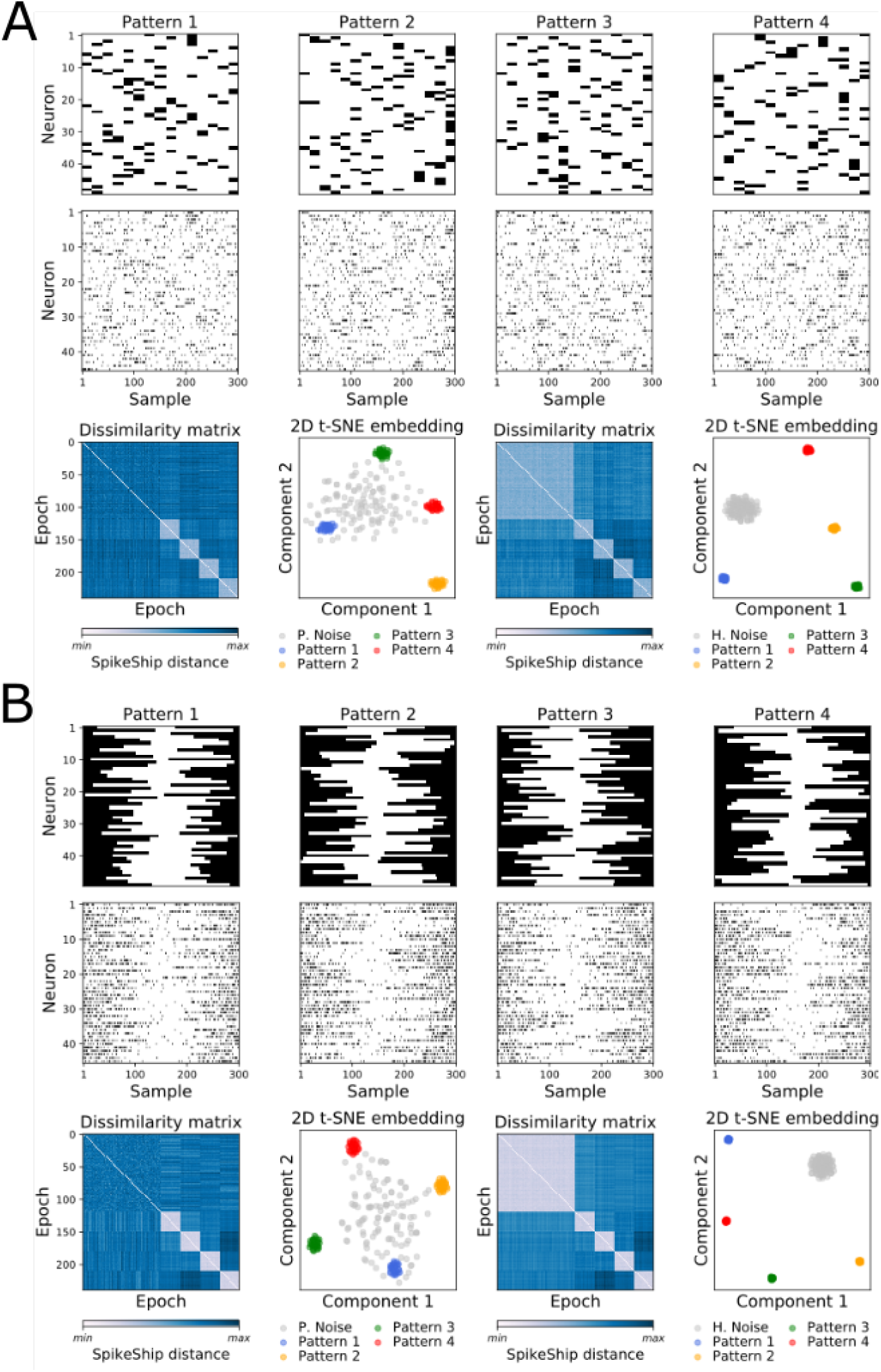
Multimodal activation and deactivation patterns can be detected using SpikeShip. Simulations from [27]. (A) Multiple bimodal activation patterns and examples of realizations for each pattern (*N* = 50 neurons). Simulation parameters were *λ*_*in*_ = 0.35 spks/sample, *λ*_*out*_ = 0.05 spks/sample, *T*_*epoch*_ = 300 and *T*_*pulse*_ = 20 samples. Bottom figures show sorted dissimilarity matrix and t-SNE for simulation with patterned noise (left) and homogeneous noise (right). (B) Multiple bimodal activation patterns and examples of realizations for each pattern (*N* = 50 neurons). Simulation parameters were *λ*_*out*_ = 0.02 spks/sample (i.e. the deactivation period), *λ*_*in*_ = 0.3 spks/sample, *T*_*epoch*_ = 300 and *T*_*deactivation*_ = 150 samples. Bottom figures show sorted dissimilarity matrix and t-SNE for simulation with patterned noise (left) and homogeneous noise (right).

**Fig S4.**
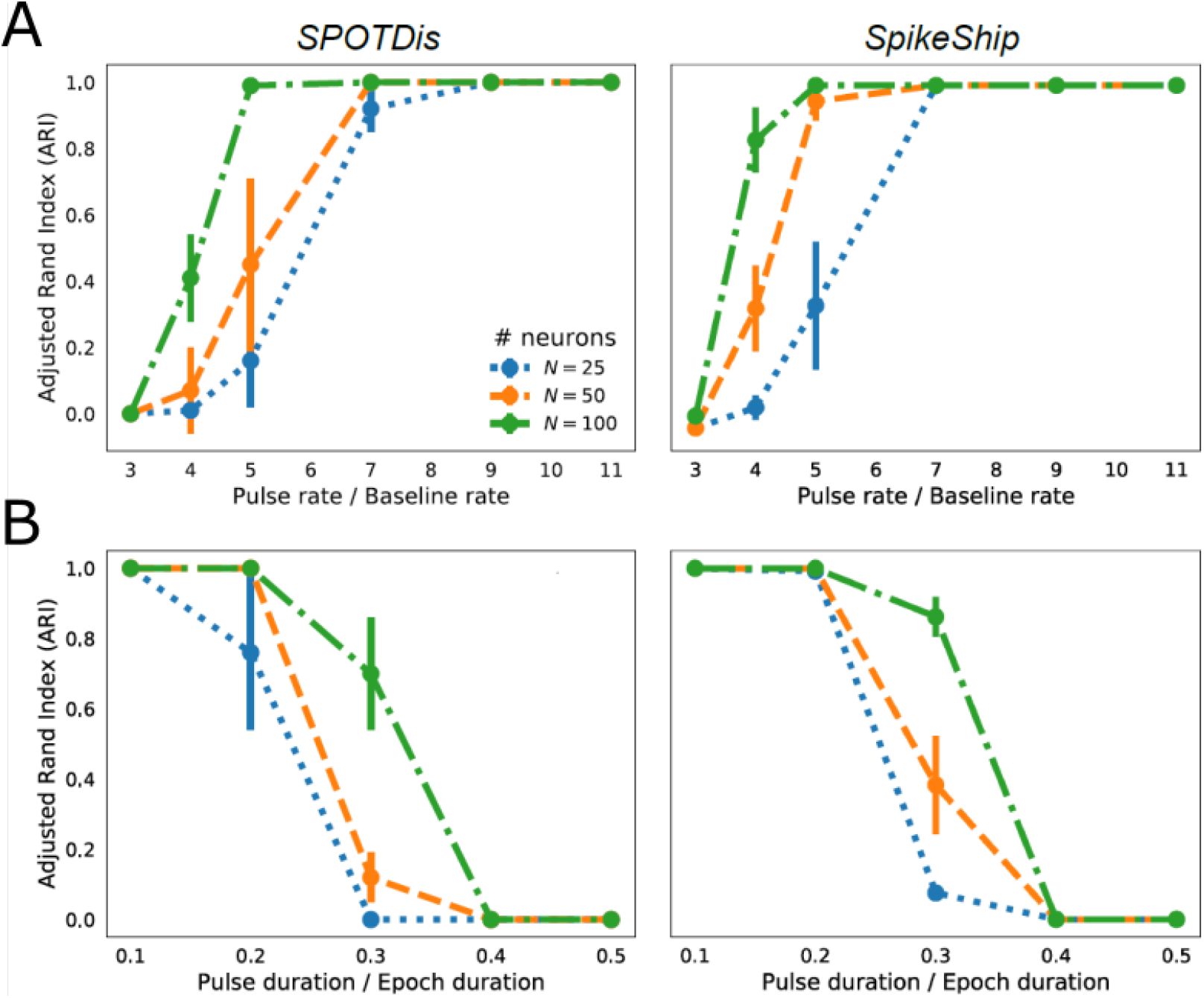
Performance of SpikeShip depends on the SNR but it outperforms SPOTDis. (A) Performance of SPOTDis (left) [27] and SpikeShip (right) measured with ARI score. We performed the same simulations as in [27]. Firing rate inside pulse period is varied, while the firing rate outside pulse was varied. We simulated 5 patterns with 30 repetitions each, with *λ*_*out*_ = 0.05 spks/sample, and *λ*_*in*_ attaining values of 0.15, 0.2, 0.25, 0.35, 0.45 or 0.5 spks/sample, *T*_*pulse*_ = 30 and *T*_*epoch*_ = 1000 samples. The number of neurons was 25, 50 or 100, and 150 epochs of homogeneous noise. We show the mean and the standard deviation across 10 repetitions of the same simulation. Performance relative to ground truth increases with SNR. Lower SNRs are needed for achieving the same performance when the number of neurons is larger. (B) as (A), but now varying the pulse duration. Simulation parameters were *λ*_*out*_ = 0.05 spks/sample, and *λ*_*in*_ = 0.5, 0.4, 0.3, 0.2, 0.1 spks/sample, and *T*_*pulse*_ of 100, 200, 300, 400 or 500 samples, with *T*_*epoch*_ = 1000 samples; note that the product of *λ*_*in*_. *T*_*pulse*_ remained constant.

**Fig S5.**
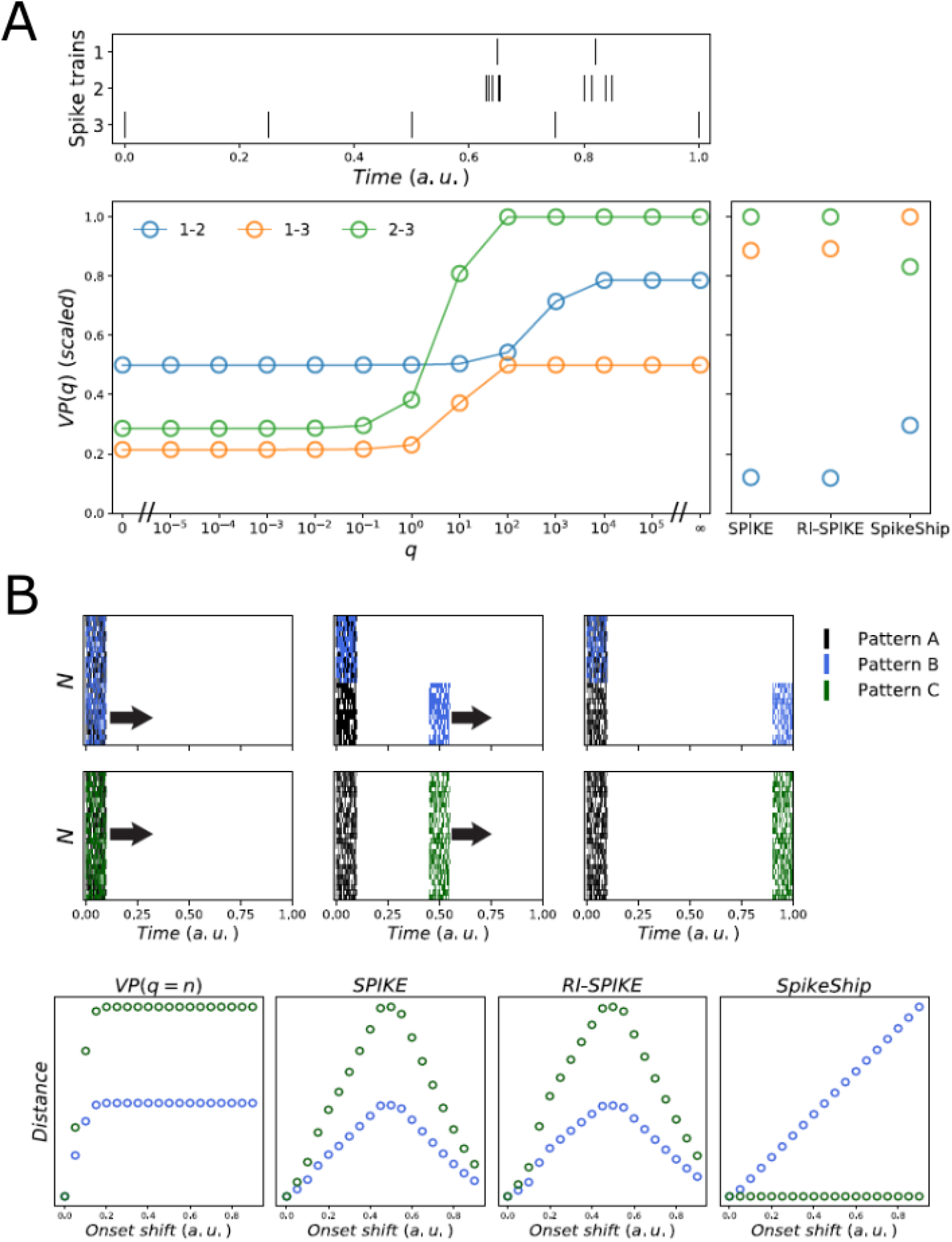
Comparison of metrics varying stimulus onset. A) SpikeShip preserves timing relationship between spike trains. Simulations of three spike trains as in [32](Fig. 5D). SpikeShip extracts the temporal information based on the timing not rate of spike trains. B) SpikeShip preserves relative timing information for multi-neuron patterns. Top: Example of three multi-neuron spiking patterns. Onset of spike patterns *B* and *C* where shifted relative to pattern *A*. Colors correspond to the shift/delay applied to half (blue) and the full neural population (green). Bottom: Comparison between spike train distances measuring patterns (*A, B*) and pattern (*A, C*). SpikeShip can detect linear changes in the neural pattern when the delay is applied to the half of the neural population. When no changes are applied in the temporal information but a global delay. SpikeShip is invariant to delays applied to the entire neural population.

**Fig S6.**
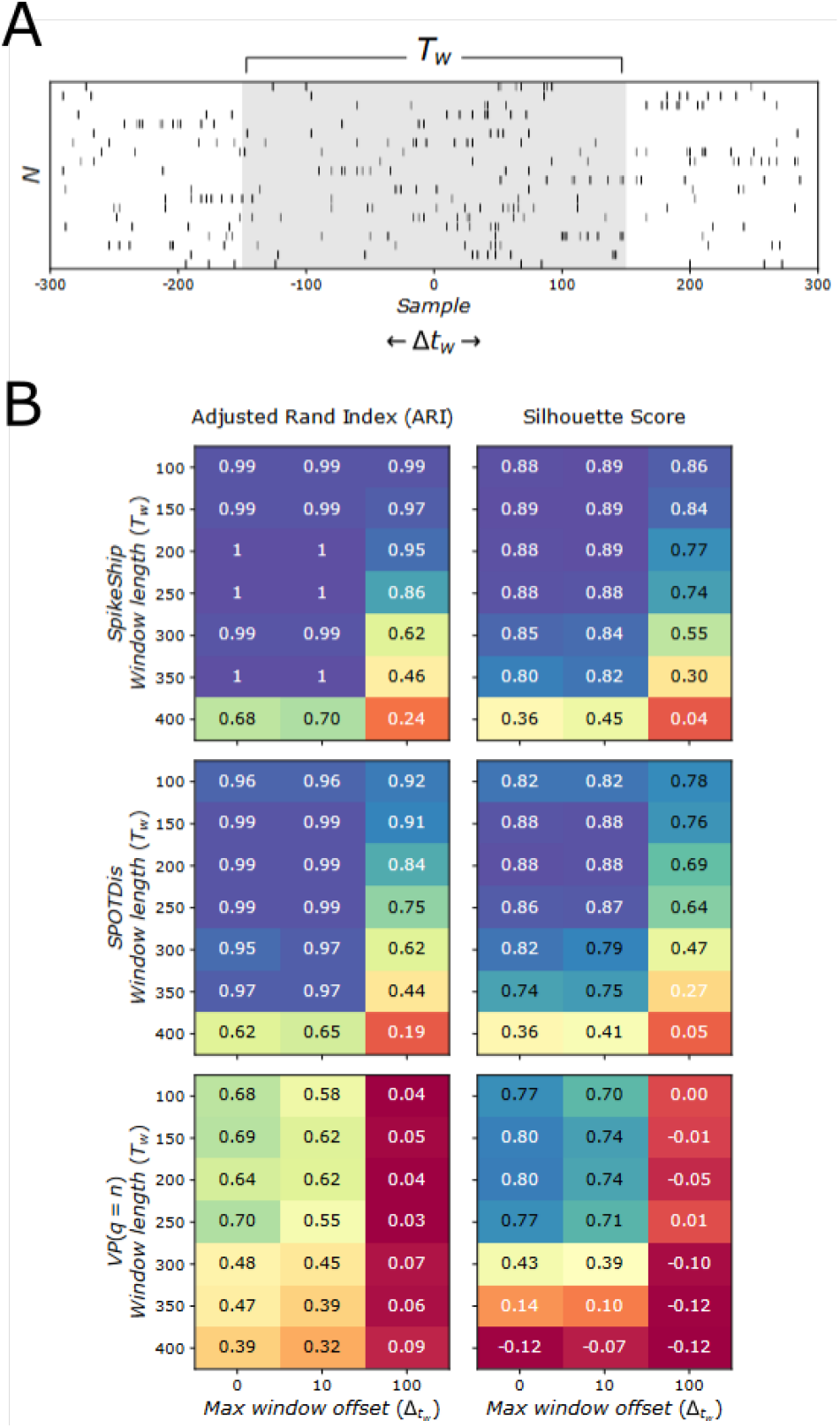
Dependence of clustering performance on chosen window length and temporal jitter of spike pattern onset. Simulations from [27]. (A) Each pattern has a length of 300 samples, and is embedded in a larger window starting from -300 samples to +300 samples, with homogeneous noise surrounding the pattern on the left and right. The onset of the pattern is -150 samples plus some random offset ∆*t*_*w*_. For each epoch realization, the value of ∆*t*_*w*_ was randomly chosen with uniform probability from an interval determined by the maximum window offset (max offset of 100 meant that ∆*t*_*w*_ *∈* [-100,100]). For the clustering, we assumed that the sequence duration is unknown. We select a window ranging from *− T*_*w*_*/*2 to +*T*_*w*_*/*2 samples of length *T*_*w*_.Clustering performance was measured relative to ground-truth (ARI) and with an unsupervised performance measure, Silhouette. Clustering performance decreased as the maximum window offset increased, due to the inclusion of noise spikes around the spike pattern. SpikeShip has a small but consistent performance advantage relative to SPOTDis. Furthermore, SpikeShip strongly outperformed VP results in clustering performance, which as expected was severely distorted by global shifts in spiking patterns. ARI and Silhouette scores correspond to the mean value obtained across 10 repetitions for each combination of window length (*T*_*w*_) and max window offset (∆*t*_*w*_).

**Fig S7.**
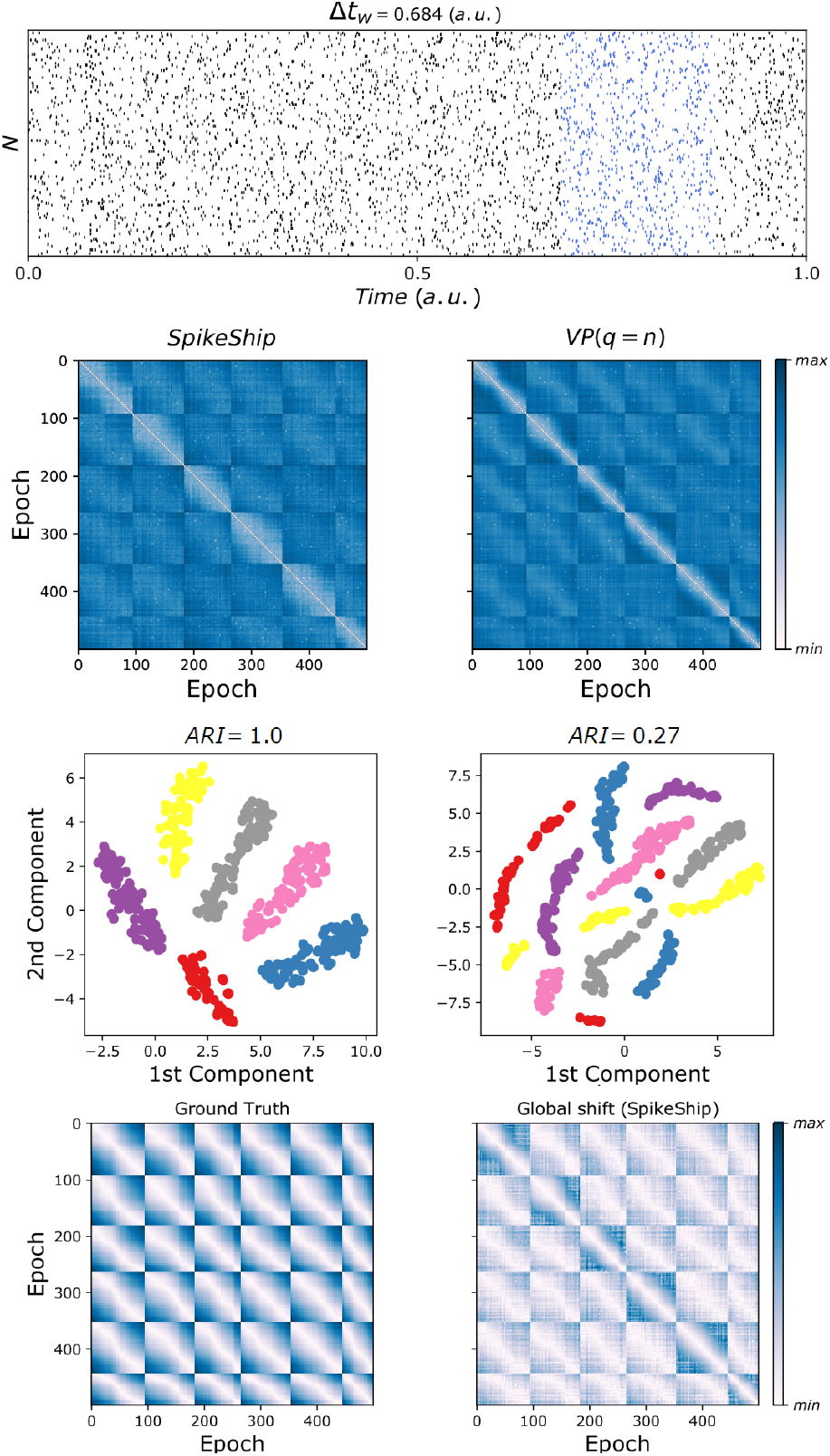
Comparison of VP and SpikeShip for simulations with multiple patterns and global shifts. Simulations from [27]. We simulated 6 patterns with Poisson noise surrounding the pattern on the left and right. The onset of the pattern was randomly assigned between 0 and 0.8 with a window length of 0.2 s for the patterns. The analysis window used here is 1 s, i.e. the entire period. SpikeShip correctly detects the 6 different patterns, but also SpikeShip can decompose the spike patterns to make it invariant to changes in global shifts. VP distance drastically depends on the global shift applied to the spikes. SpikeShip can retrieve the global shift from the spike sequences and reconstruct the random global shifts applied to the spike trains.

**Fig S8.**
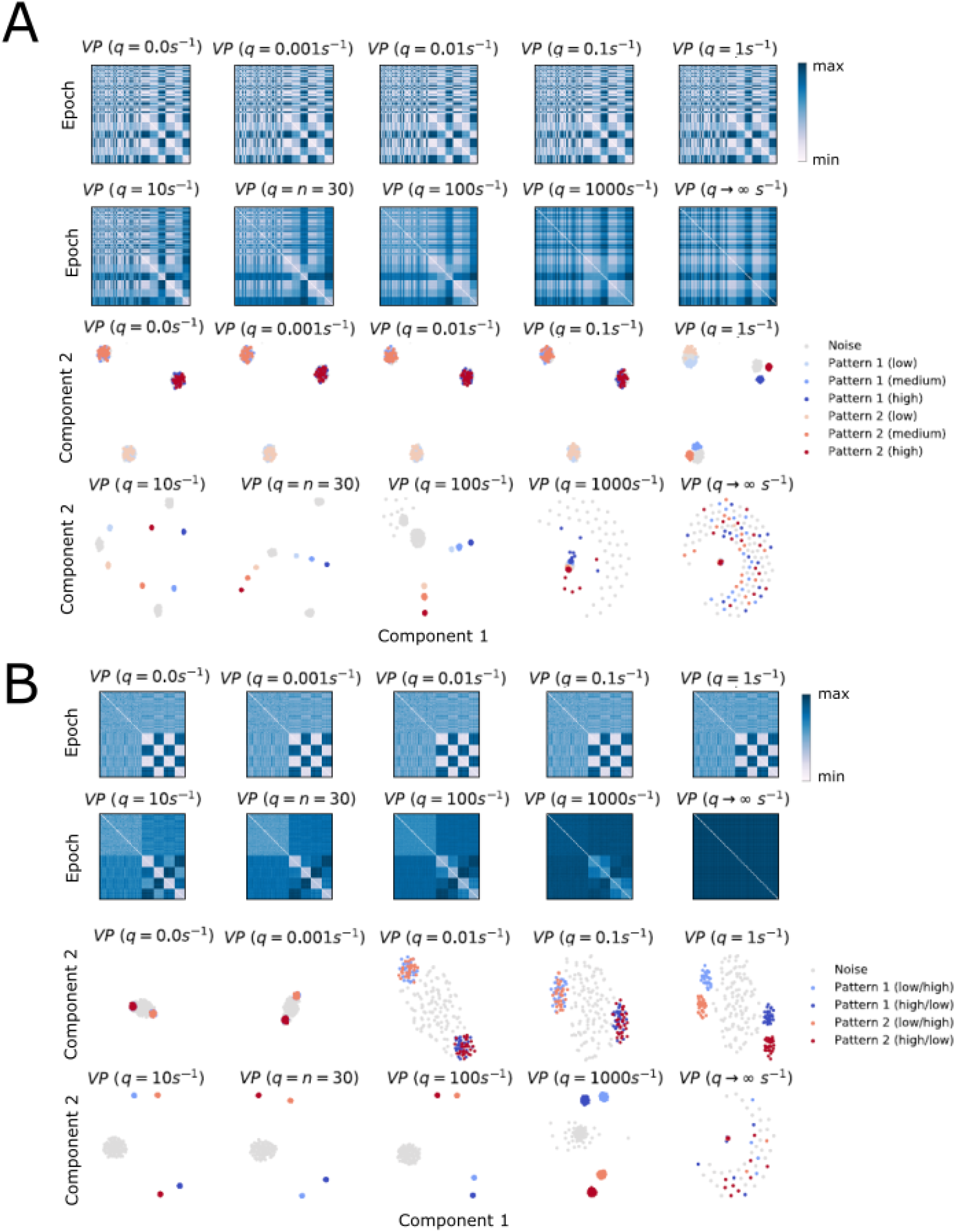
Performance of VP distance is affected by changes in both local and global scaling rates. Simulations from [27]. A) Global scaling. Same simulations as in Fig.S9. Victor-Purpura distance (VP) was used with different values of *q*. When *q* = 0, *V P* = | *n*_*i*_ *−n*_*j*_|, with *n*_*i*_ and *n*_*j*_ the spike count of spike sequences *i* and *j*, respectively. Epochs are clustered based on rates. B) Local scaling. Same simulations as in Fig.S10. VP distance was used with different values of *q*. When *q → ∞, V P* = *n*_*i*_ + *n*_*j*_. Besides high values of *q* aim to extract temporal information from spike trains, these 2D embeddings demonstrate that the contribution between rate and timing using VP is difficult to interpret and very sensitive to noise.

**Fig S9.**
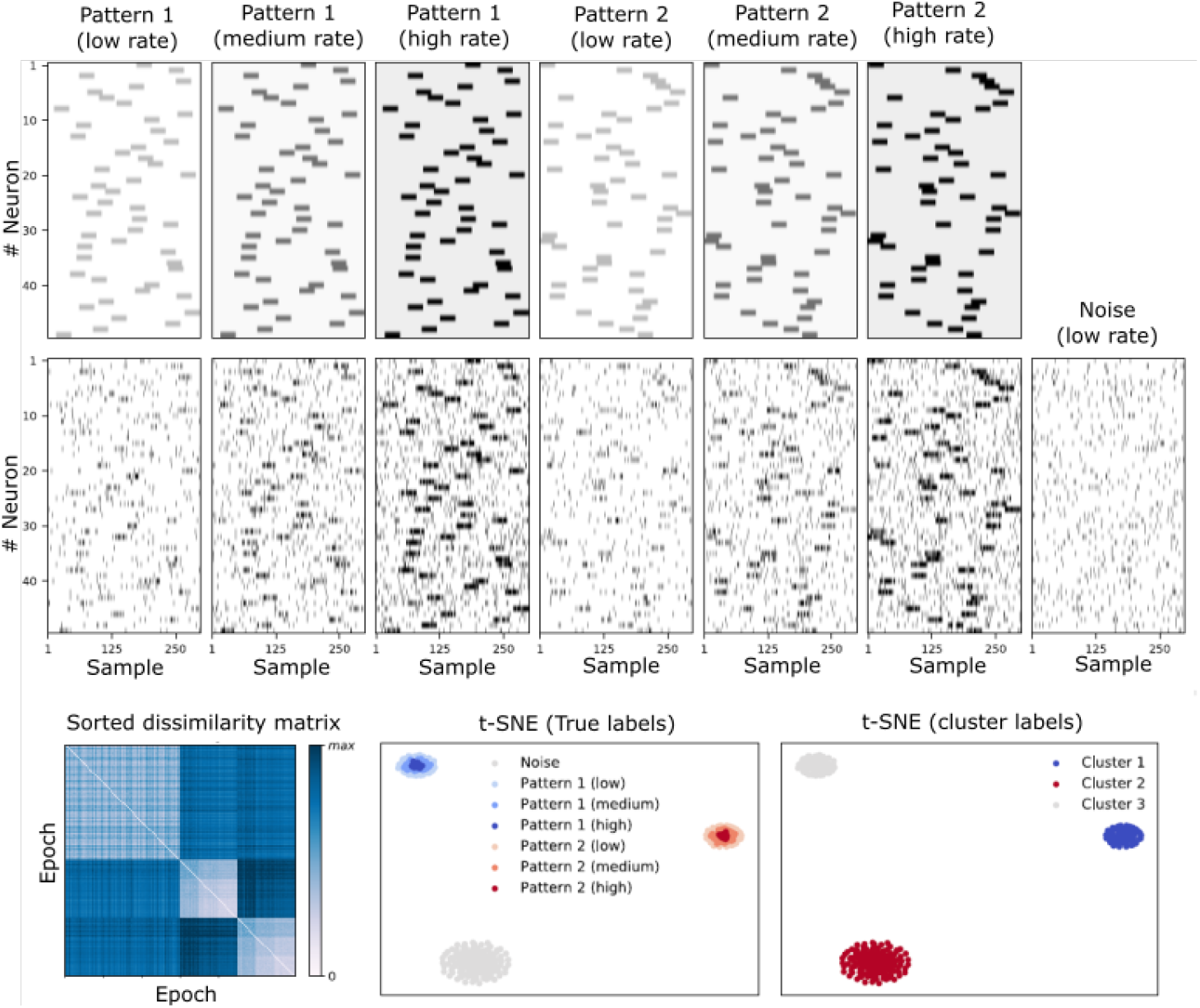
Performance SpikeShip is not affected by a global scaling rate. Simulations from [27]. Shown are two different temporal patterns. Each temporal pattern can occur in a low (*λ*_*in*_ = 0.2 and *λ*_*out*_ = 0.02 spks/sample), medium (*λ*_*in*_0.4 and *λ*_*out*_ = 0.04 spks/sample) or high rate (*λ*_*in*_ = 0.7 and *λ*_*out*_0.07 spks/sample) state, with a constant ratio of *λ*_*in*_*/λ*_*out*_. In addition, the noise pattern can also occur in one of three rate states. The pulse duration was 30 samples. Shown at the bottom the sorted dissimilarity matrix with SpikeShip values, the t-SNE embedding with the ground-truth cluster labels and the t-SNE embedding with the HDBSCAN cluster labels.

**Fig S10.**
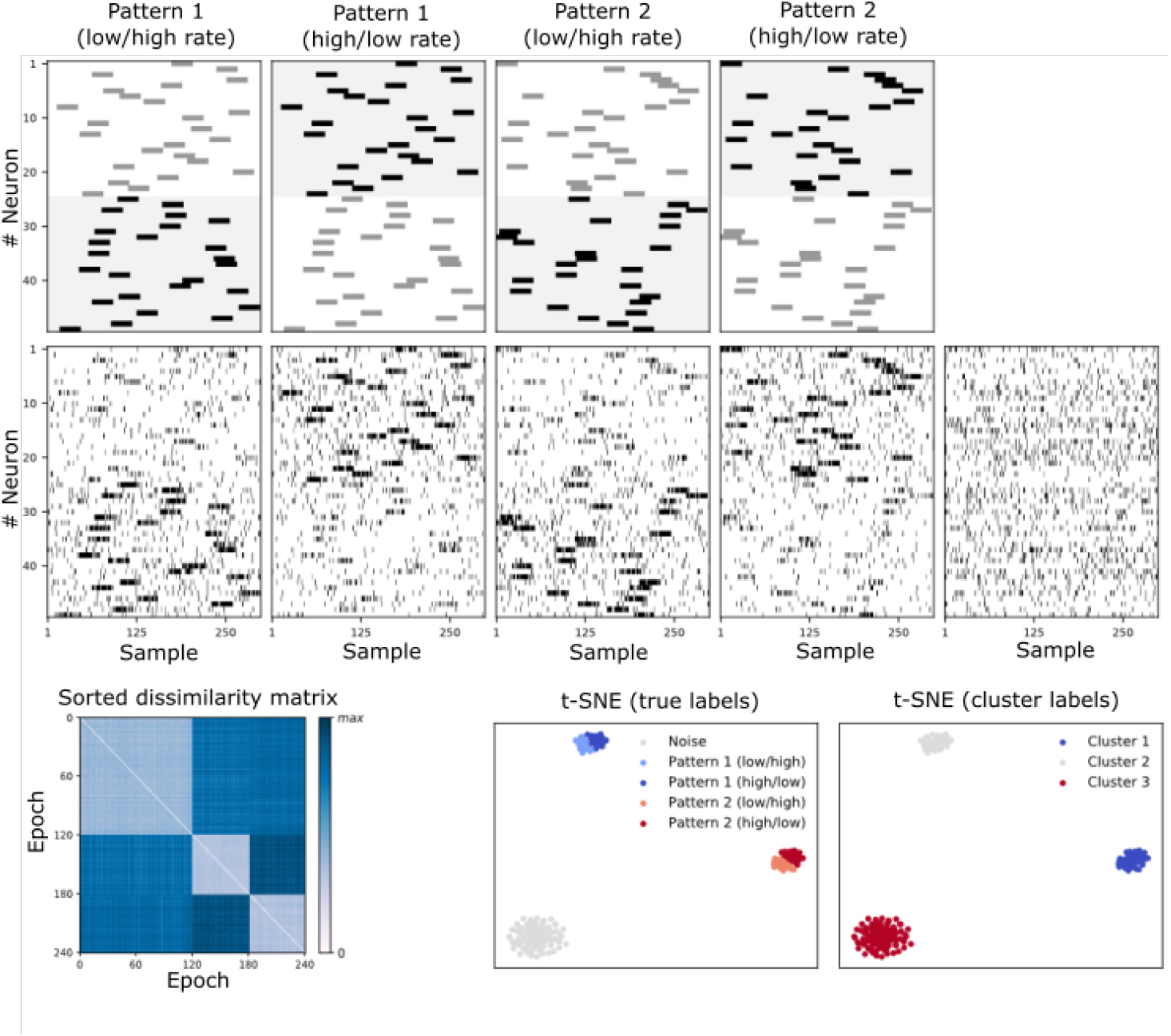
Performance of SpikeShip is not affected by a local scaling rate. Simulations from [27]. Shown are two temporal patterns. Each temporal pattern could occur in one of two rates states: In the first rate state, the first 25 neurons are firing at a low rate (*λ*_*in*_ = 0.3 and *λ*_*out*_ = 0.03 spks/sample), and the other 25 are firing at a high rate (*λ*_*in*_ = 0.7 and *λ*_*out*_ = 0.07 spks/sample). In the second rate state, the rate scaling is reversed. The pulse duration was 30 samples. Shown at the bottom the sorted dissimilarity matrix with SpikeShip values, the t-SNE embedding with the ground-truth cluster labels and the t-SNE embedding with the HDBSCAN cluster labels.

**Fig S11.**
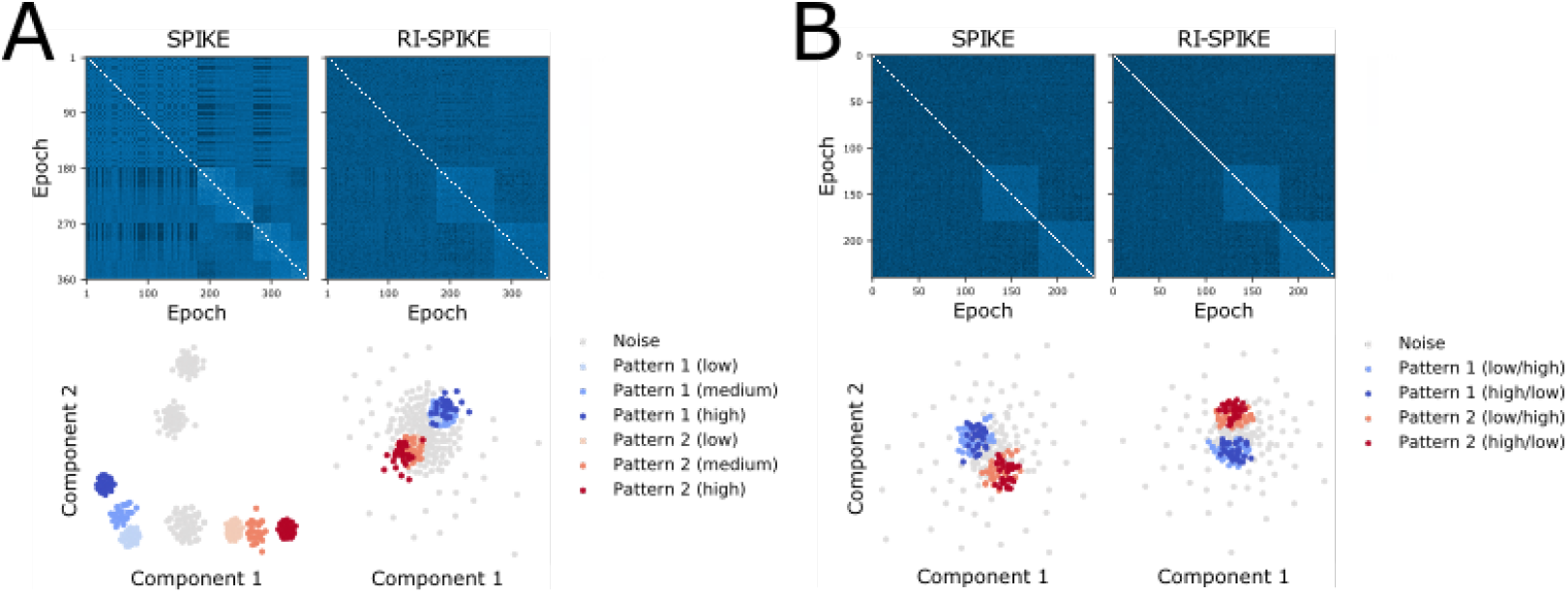
Performance of SPIKE and RI-SPIKE are affected by both global and local scaling. Simulations from [27]. A) Global scaling. SPIKE and RI-SPIKE computations for globally scaled sequences. Top: dissimilarity matrices sorted by pattern id and scaling factor. Bottom: 2D t-SNE embeddings of epochs. B) Local scaling. SPIKE and RI-SPIKE computations for locally scaled sequences. Both A and B were computed using the same simulations as in Fig. S9. Top: dissimilarity matrices sorted by pattern id and scaling factor. Bottom: 2D t-SNE embeddings of epochs. The first 180 epochs correspond to noise.

**Fig S12.**
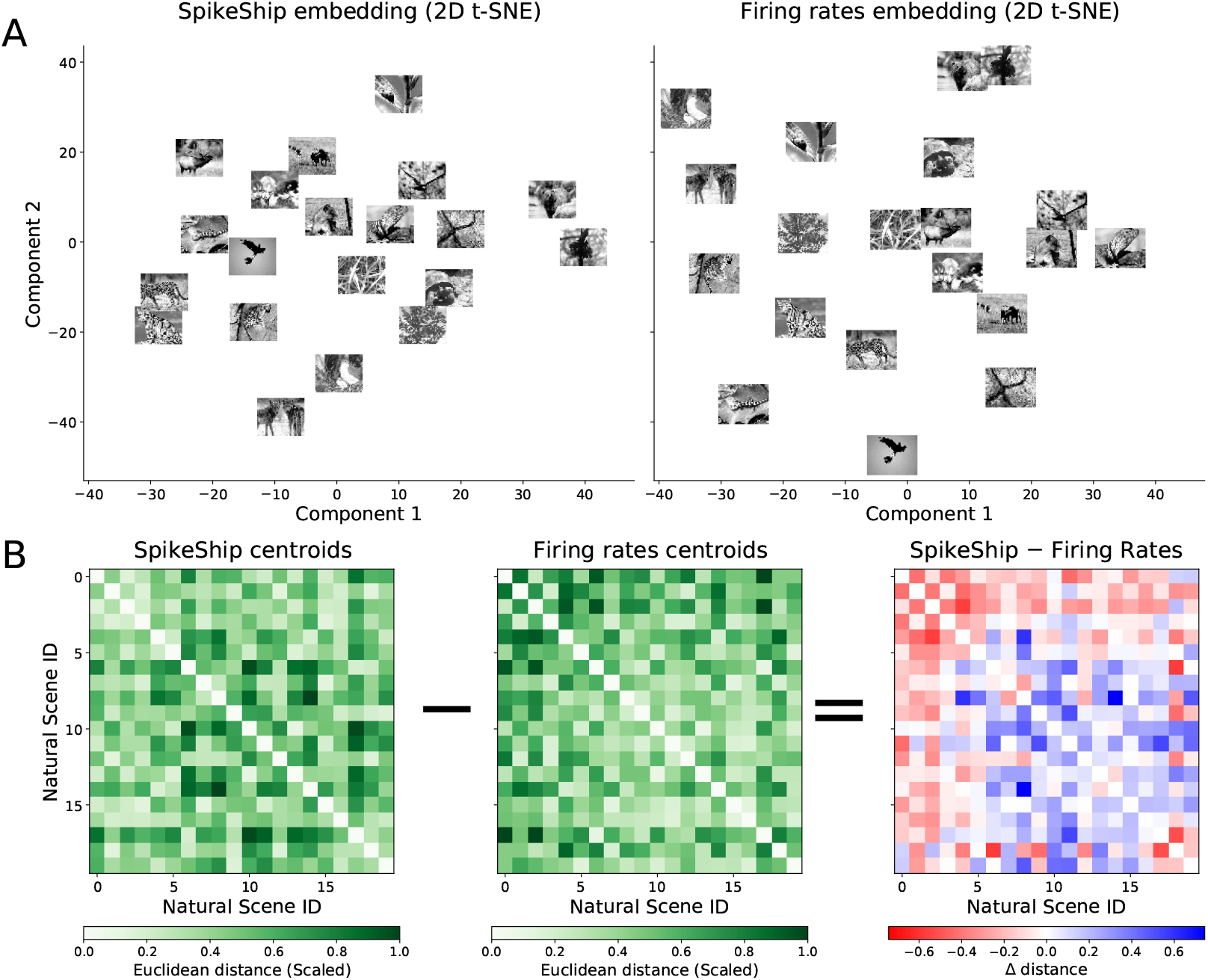
Comparison between clusters from SpikeShip and Firing rates embeddings of Natural scenes. A) 2D t-SNE embeddings from SpikeShip and Firing rates’ dissimilarity matrices. The allocation of natural scenes’ clusters are different between the two embeddings. B) Scaled Euclidean pairwise distance between centroids of each cluster for both SpikeShip (Left) and firing rates (Middle), and their difference (Right).

**Fig S13.**
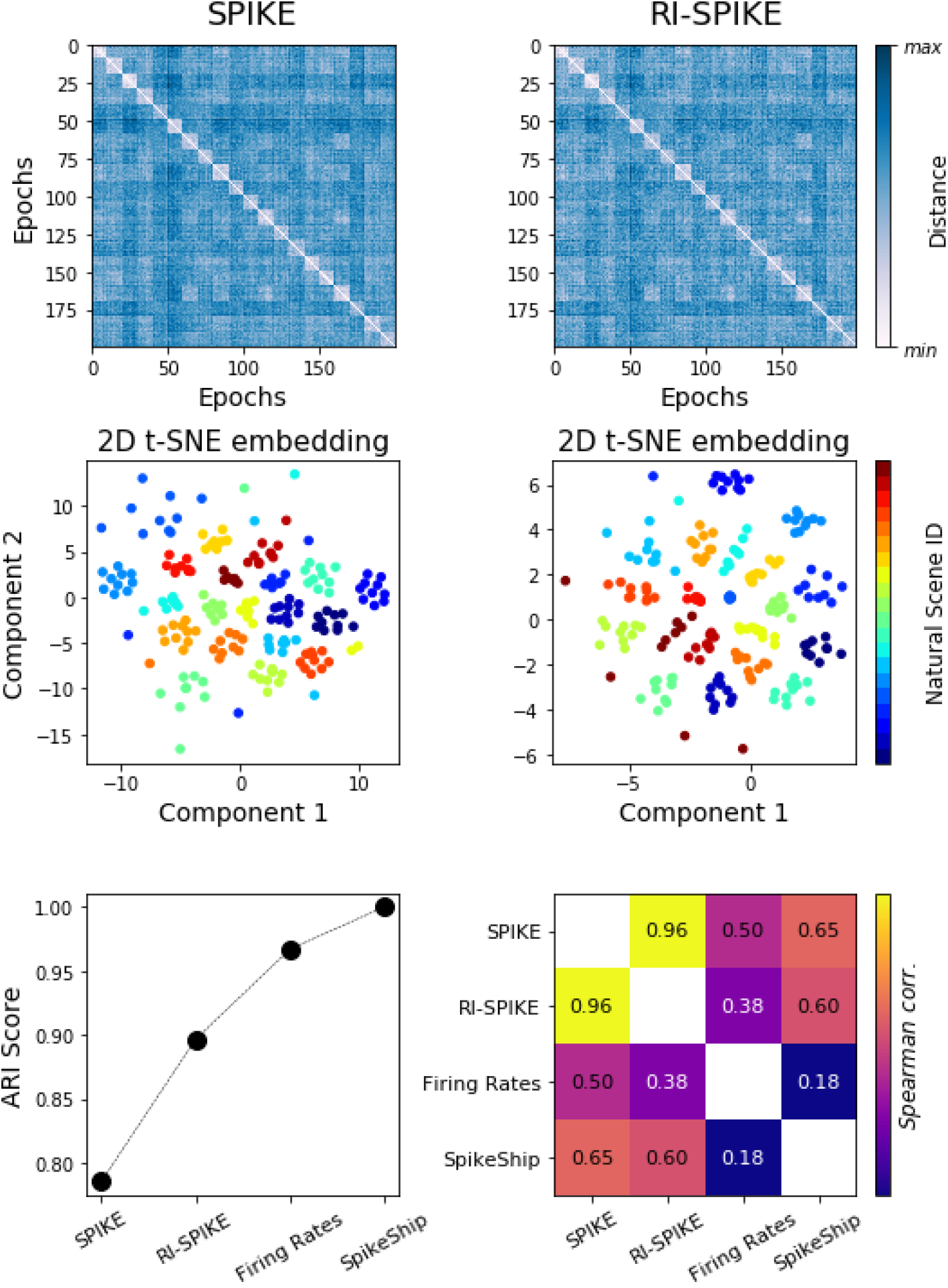
Analysis of large scale neural recordings during visual stimuli presentations with SPIKE and RI-SPIKE. Top: Dissimilarity matrices sorted by Natural Scene ID. Middle: 2D t-SNE embeddings from dissimilarity matrices colored by Natural Scene ID. Bottom: The clustering performance through ARI score and Spearman correlation between dissimilarity matrices computed via SPIKE, RI-SPIKE, Firing rates, and SpikeShip. The clustering performance of SPIKE and RI-SPIKE is lower than the clustering performance by using the traditional firing rates and SpikeShip. SPIKE and RI-SPIKE are highly correlated while Firing Rates and SpikeShip are highly uncorrelated.

**Fig S14.**
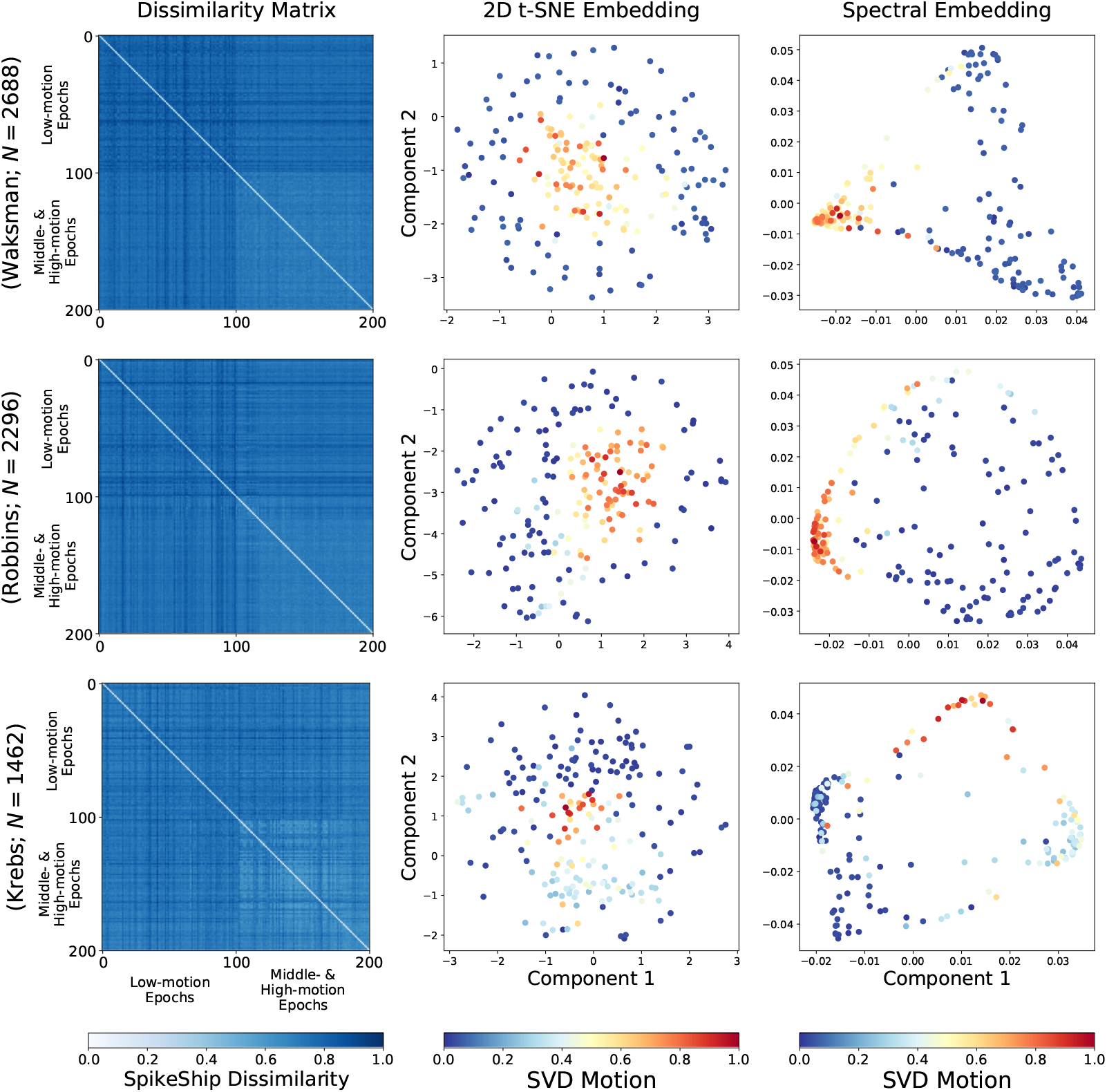
Spontaneous activity analyses for 3 mice. Multi-spike sequence analyses for three mice (rows). Left: dissimilarity matrices. Middle: 2D t-SNE embedding. Right: 2D Spectral Embedding (Laplacian Eigenmaps).

## References

1. Havenith MN, Yu S, Biederlack J, Chen NH, Singer W, Nikolic D. Synchrony makes neurons fire in sequence, and stimulus properties determine who is ahead. J Neurosci. 2011;31(23):8570–84.

2. Vinck M, Lima B, Womelsdorf T, 𝒪ostenveld R, Singer W, Neuenschwander S, et al. Gamma-phase shifting in awake monkey visual cortex. Journal of Neuroscience. 2010;30(4):1250–1257.

3. Luczak A, Bartho P, Harris KD. Spontaneous events outline the realm of possible sensory responses in neocortical populations. Neuron. 2009;62(3):413–25.

4. Lu T, Liang L, Wang X. Temporal and rate representations of time-varying signals in the auditory cortex of awake primates. Nat Neurosci. 2001;4(11):1131–8.

5. Pastalkova E, Itskov V, Amarasingham A, Buzsaki G. Internally generated cell assembly sequences in the rat hippocampus. Science. 2008;321(5894):1322–7.

6. Ikegaya Y, Aaron G, Cossart R, Aronov D, Lampl I, Ferster D, et al. Synfire chains and cortical songs: temporal modules of cortical activity. Science. 2004;304(5670):559–564.

7. Wehr M, Laurent G. 𝒪dour encoding by temporal sequences of firing in oscillating neural assemblies. Nature. 1996;384(6605):162–6.

8. Seung HS, Lee DD. The manifold ways of perception. science. 2000;290(5500):2268– 2269.

9. Jun JJ, Steinmetz NA, Siegle JH, Denman DJ, Bauza M, Barbarits B, et al. Fully integrated silicon probes for high-density recording of neural activity. Nature. 2017;551(7679):232–236.

10. Stringer C, Pachitariu M, Steinmetz N, Reddy CB, Carandini M, Harris KD. Spontaneous behaviors drive multidimensional, brainwide activity. Science. 2019;364(6437).

11. Stringer C, Michaelos M, Tsyboulski D, Lindo SE, Pachitariu M. High-precision coding in visual cortex. Cell. 2021;184(10):2767–2778.

12. Siegle JH, Jia X, Durand S, Gale S, Bennett C, Graddis N, et al. Survey of spiking in the mouse visual system reveals functional hierarchy. Nature. 2021;592(7852):86– 92.

13. Chang L, Tsao DY. The code for facial identity in the primate brain. Cell. 2017;169(6):1013–1028.

14. Churchland MM, Cunningham JP, Kaufman MT, Foster JD, Nuyujukian P, Ryu SI, et al. Neural population dynamics during reaching. Nature. 2012;487(7405):51–56.

15. Dragoi G, Buzsaki G. Temporal encoding of place sequences by hippocampal cell assemblies. Neuron. 2006;50(1):145–57.

16. Mainen ZF, Sejnowski TJ. Reliability of spike timing in neocortical neurons. Science. 1995;268(5216):1503–1506.

17. Markram H, Lubke J, Frotscher M, Sakmann B. Regulation of synaptic efficacy by coincidence of postsynaptic APs and EPSPs. Science. 1997;275(5297):213–5.

18. Dan Y, Poo MM. Spike timing-dependent plasticity of neural circuits. Neuron. 2004;44(1):23–30.

19. Abbott LF, Blum KI. Functional significance of long-term potentiation for sequence learning and prediction. Cereb Cortex. 1996;6(3):406–16.

20. Buonomano DV, Maass W. State-dependent computations: spatiotemporal processing in cortical networks. Nat Rev Neurosci. 2009;10(2):113–25.

21. Singer W, Lazar A. Does the cerebral cortex exploit high-dimensional, non-linear dynamics for information processing? Frontiers in computational neuroscience. 2016;10:99.

22. Steinmetz N, Pachitariu M, Stringer C, Carandini M, Harris K. Eight-probe Neuropixels recordings during spontaneous behaviors; 2019. Available from: https://janelia.figshare.com/articles/dataset/Eight-probe_Neuropixels_recordings_during_spontaneous_behaviors/7739750/4.

23. Victor JD, Purpura KP. Nature and precision of temporal coding in visual cortex: a metric-space analysis. Journal of neurophysiology. 1996;76(2):1310–1326.

24. Victor JD, Purpura KP. Metric-space analysis of spike trains: theory, algorithms and application. Network: computation in neural systems. 1997;8(2):127–164.

25. Kreuz T, Chicharro D, Houghton C, Andrzejak RG, Mormann F. Monitoring spike train synchrony. Journal of neurophysiology. 2013;109(5):1457–1472.

26. Satuvuori E, Mulansky M, Bozanic N, Malvestio I, Zeldenrust F, Lenk K, et al. Measures of spike train synchrony for data with multiple time scales. Journal of neuroscience methods. 2017;287:25–38.

27. Grossberger L, Battaglia FP, Vinck M. Unsupervised clustering of temporal patterns in high-dimensional neuronal ensembles using a novel dissimilarity measure. PLoS computational biology. 2018;14(7):e1006283.

28. Van der Maaten L, Hinton G. Visualizing data using t-SNE. Journal of machine learning research. 2008;9(11).

29. Van Der Maaten L. Accelerating t-SNE using tree-based algorithms. The Journal of Machine Learning Research. 2014;15(1):3221–3245.

30. Hinton G. Stochastic neighbor embedding. Advances in neural information processing systems. 2003;15:857–864.

31. McInnes L, Healy J, Astels S. hdbscan: Hierarchical density based clustering. The Journal of 𝒪pen Source Software. 2017;2(11).

32. Satuvuori E, Kreuz T. Which spike train distance is most suitable for distinguishing rate and temporal coding? Journal of neuroscience methods. 2018;299:22–33.

33. Thorpe S, Delorme A, Van Rullen R. Spike-based strategies for rapid processing. Neural networks. 2001;14(6-7):715–725.

34. McGinley MJ, Vinck M, Reimer J, Batista-Brito R, Zagha E, Cadwell CR, et al. Waking State: Rapid Variations Modulate Neural and Behavioral Responses. Neuron. 2015;87(6):1143–61.

35. Williams AH, Poole B, Maheswaranathan N, Dhawale AK, Fisher T, Wilson CD, et al. Discovering precise temporal patterns in large-scale neural recordings through robust and interpretable time warping. Neuron. 2020;105(2):246–259.

36. Sihn D, Kim SP. A spike train distance robust to firing rate changes based on the Earth Mover’s Distance. Frontiers in Computational Neuroscience. 2019;13:82.

37. Cormen TH, Leiserson CE, Rivest RL, Stein C. Introduction to algorithms. MIT press; 2009.

38. Bleich C, 𝒪verton ML. A linear-time algorithm for the weighted median problem. Courant Institute of Mathematical Sciences, New York University; 1983.

39. Bovo F. Robustats; 2020. https://github.com/FilippoBovo/robustats.

40. Denker M, Yegenoglu A, Grün S. Collaborative HPC-enabled workflows on the HBP Collaboratory using the Elephant framework. In: Neuroinformatics 2018; 2018. p. P19. Available from: https://abstracts.g-node.org/conference/NI2018/abstracts#/uuid/023bec4e-0c35-4563-81ce-2c6fac282abd.

41. Mulansky M, Kreuz T. PySpike—A Python library for analyzing spike train synchrony. SoftwareX. 2016;5:183–189.

